# Deep learning from phylogenies to uncover the epidemiological dynamics of outbreaks

**DOI:** 10.1101/2021.03.11.435006

**Authors:** J Voznica, A Zhukova, V Boskova, E Saulnier, F Lemoine, M Moslonka-Lefebvre, O Gascuel

**Affiliations:** Institut Pasteur, Université Paris Cité, Unité Bioinformatique Evolutive, Paris, FRANCE; Université de Paris, Paris, FRANCE; Institut de Biologie de l’École Normale Supérieure, Ecole Normale Supérieure, CNRS, INSERM, Université Paris Sciences et Lettres, Paris, FRANCE; Institut Pasteur, Université Paris Cité, Bioinformatics and Biostatistics Hub, Paris, FRANCE; Institut Pasteur, Université Paris Cité, Epidemiology and Modelling of Antibiotic Evasion, Paris, FRANCE; Université Paris-Saclay, UVSQ, Inserm, CESP, Villejuif, FRANCE; Center for Integrative Bioinformatics Vienna, Max Perutz Labs, University of Vienna and Medical University of Vienna, Vienna, AUSTRIA; Institut de Systématique, Evolution, Biodiversité (UMR 7205 - CNRS, Muséum National d’Histoire Naturelle, SU, EPHE, UA), Paris, FRANCE

**Keywords:** Phylodynamics, molecular epidemiology, tree representation, neural networks, HIV.

## Abstract

Widely applicable, accurate and fast inference methods in phylodynamics are needed to fully profit from the richness of genetic data in uncovering the dynamics of epidemics. Standard methods, including maximum-likelihood and Bayesian approaches, generally rely on complex mathematical formulae and approximations, and do not scale with dataset size. We develop a likelihood-free, simulation-based approach, which combines deep learning with (1) a large set of summary statistics measured on phylogenies or (2) a complete and compact representation of trees, which avoids potential limitations of summary statistics and applies to any phylodynamics model. Our method enables both model selection and estimation of epidemiological parameters from very large phylogenies. We demonstrate its speed and accuracy on simulated data, where it performs better than the state-of-the-art methods. To illustrate its applicability, we assess the dynamics induced by superspreading individuals in an HIV dataset of men-having-sex-with-men in Zurich. Our tool PhyloDeep is available on github.com/evolbioinfo/phylodeep.

## INTRODUCTION

Pathogen phylodynamics is a field combining phylogenetics and epidemiology^[1]^. Viral or bacterial samples from patients are sequenced and used to infer a phylogeny, which describes the pathogen’s spread among patients. The tips of such phylogenies represent sampled pathogens, and the internal nodes transmission events. Moreover, transmission events can be dated and thereby provide hints on transmission patterns. Such information is extracted by phylodynamic methods to estimate epidemiological and population dynamic parameters^[2–4]^, assess the impact of population structure^[2, 5]^, and reveal the origins of epidemics^[6]^.

Birth-death models^[7]^ incorporate easily interpretable parameters common to standard infectious-disease epidemiology, such as basic reproduction number R_0_, infectious period, *etc.* In contrast to the standard epidemiological models, the birth-death models can be applied to estimate parameters from phylogenetic trees^[8]^. In these models, births represent transmission events, while deaths represent removal events for example due to treatment or recovery. Upon a patient’s removal, their pathogens can be sampled, producing tips in the tree.

Here we focus on three specific, well-established birth-death models (**Fig. 1)**: birth-death model (BD)^[8, 9]^, birth-death model with exposed and infectious classes (BDEI)^[5,10,11]^, and birth-death model with superspreading (BDSS)^[5, 12]^. These models were deployed using BEAST2^[12, 13]^ to study the phylodynamics of such diverse pathogens as Ebola virus^[10]^, Influenza virus^[12]^, Human Immunodeficiency Virus (HIV)^[5]^, Zika^[14]^ or SARS-CoV-2^[15]^. Using these models, we will demonstrate the reliability of our deep learning-based approach.

**Fig. 1:**
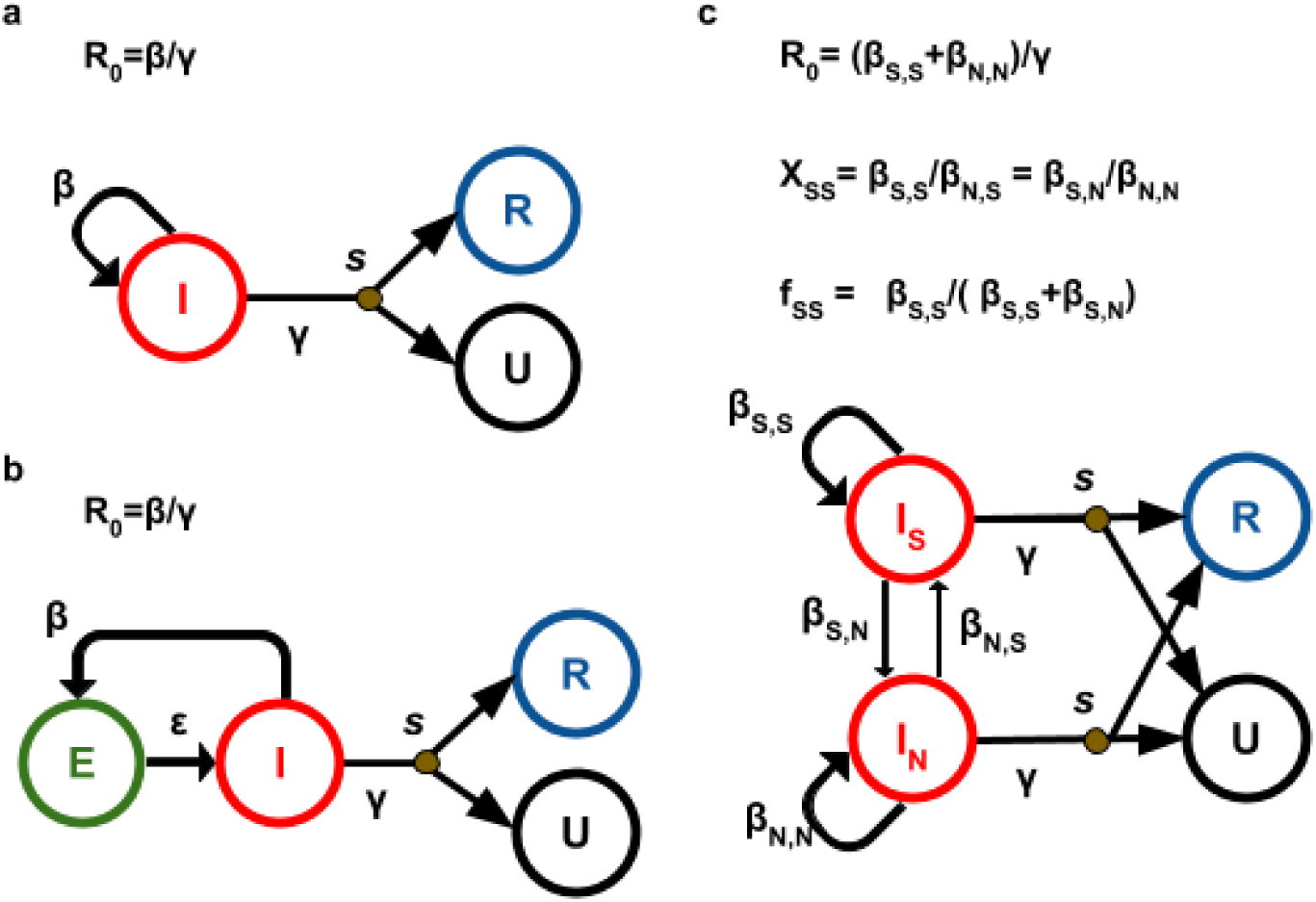
Birth-death models. Note to Fig. 1. **a** Birth-death model (BD)^[8, 9]^, **b,** birth-death model with Exposed-Infectious individuals (BDEI)^[5,10,11]^ and **c,** birth-death model with SuperSpreading (BDSS)^[5, 12]^. BD is the simplest generative model, used to estimate R0 and the infectious period (1/γ)^[8, 9]^. BDEI and BDSS are extended version of BD. BDEI enables to estimate latency period (1/ε) during which individuals of exposed class E are infected, but not infectious^[5,10,11]^. BDSS includes two populations with heterogeneous infectiousness: the so-called superspreading individuals (S) and normal spreaders (N). Superspreading individuals are present only at a low fraction in the population (f_ss_) and may transmit the disease at a rate that is multiple times higher than that of normal spreaders (rate ratio = X_ss_)^[5, 12]^. Superspreading can have various complex causes, such as the heterogeneity of immune response, disease progression, co-infection with other diseases, social contact patterns or risk behaviour, *etc*. Infectious individuals I (superspreading infectious individuals I_S_ and normal spreaders I_N_ for BDSS), transmit the disease at rate β (β_X,Y_ for an individual of type X transmitting to an individual of type Y for BDSS), giving rise to a newly infected individual. The newly infected individual is either infectious right away in BD and BDSS or goes through an exposed state before becoming infectious at rate ε in BDEI. Infectious individuals are removed at rate γ. Upon removal, they can be sampled with probability *s*, becoming of removed sampled class R. If not sampled upon removal, they move to non-infectious unsampled class U.

While a great effort has been invested in the development of new epidemiological models in phylodynamics, the field has been slowed down by the mathematical complexity inherent to these models. BD, the simplest model, has a closed form solution for the likelihood formula of a tree for a given set of parameters^[8, 10]^, but more complex models (*e.g.,* BDEI and BDSS) rely on a set of ordinary differential equations (ODEs) that cannot be solved analytically. To estimate parameter values through maximum-likelihood and Bayesian approaches, these ODEs must be approximated numerically for each tree node^[5,10–12]^. These calculations become difficult as the tree size increases, resulting in numerical instability and inaccuracy^[12]^, as we will see below.

Inference issues with complex models are typically overcome by approximate Bayesian computation (ABC)^[16, 17]^. ABC is a simulation-based technique relying on a rejection algorithm^[18]^, where from a set of simulated phylogenies within a given prior (values assumed for parameter values), those closest to the analysed phylogeny are retained and give the posterior distribution of the parameters. This scheme relies on the definition of a set of summary statistics aimed at representing a phylogeny and on a distance measure between trees. This approach is thus sensitive to the choice of the summary statistics and distance function (*e.g.,* Euclidean distance). To address this issue Saulnier *et al.*^[19]^ developed a large set of summary statistics. In addition, they used a regression step to select the most relevant statistics and to correct for the discrepancy between the simulations retained in the rejection step and the analysed phylogeny. They observed that the sensitivity to the rejection parameters were greatly attenuated thanks to regression (see also Blum *et al.*^[20]^).

Our work is a continuation of regression-based ABC, and aims at overcoming its main limitations. Using the approximation power of currently available neural network architectures, we propose a likelihood-free method relying on deep learning from millions of trees of varying size simulated within a broad range of parameter values. By doing so, we bypass the rejection step, which is both time consuming with large simulation sets, and sensitive to the choice of the distance function and summary statistics. To describe simulated trees and use them as input for the deep learner, we develop two tree representations: (1) a large set of summary statistics mostly based on Saulnier *et al.*^[19]^, and (2) a complete and compact vectorial representation of phylogenies, including both the tree topology and branch lengths. The summary statistics are derived from our understanding and knowledge of the epidemiological processes. However, they can be incomplete and thus miss some important aspects of the studied phylogenies, which can potentially result in low accuracy during inference. Moreover, it is expected that new phylodynamic models will require design of new summary statistics, as confirmed by our results with BDSS. In contrast, our vectorial representation is a raw data representation that preserves all information contained in the phylogeny and thus should be accurate and deployable on any new model, provided the model parameters are identifiable. Our vectorial representation naturally fits with deep learning methods, especially the convolutional architectures, which have already proven their ability to extract relevant features from raw representations, for example in image analysis^[21, 22]^ or weather prediction^[23]^.

In the following, we introduce our vectorial tree representation and the new summary statistics designed for BDSS. We then present the deep learning architectures trained on these representations and evaluate their accuracy on simulated datasets in terms of both parameter estimation and model selection. We show that our approach applies not only to trees of the same size as the training instances, but also to very large trees with thousands of tips through the analysis of their subtrees. The results are compared to those of the gold standard method, BEAST2^[12, 13]^. Lastly, we showcase our methods on an HIV dataset^[24, 25]^ from the men-having-sex-with-men (MSM) community from Zurich. All technical details are provided in **Methods**. Our methods and tools are implemented in the PhyloDeep software, which is available on GitHub (<u>github.com/evolbioinfo/phylodeep</u>), PyPi (<u>pypi.org/project/phylodeep</u>) and Docker Hub (<u>hub.docker.com/r/evolbioinfo/phylodeep</u>).

## RESULTS

Neural networks are trained on numerical vectors from which they can learn regression and classification tasks. We trained such networks on phylogenetic trees to estimate epidemiological parameters (regression) and select phylodynamic models (classification). We undertook two strategies for representing phylogenetic trees as numerical vectors, which we describe first, before showing the results with simulated and real data.

### Summary statistics (SS) representation

We used a set of 83 SS developed by Saulnier *et al.*^[19]^: 26 measures of branch lengths, such as median of both internal and tip branch lengths; 8 measures of tree topology, such as tree imbalance; 9 measures on the number of lineages through time, such as time and height of its maximum; and 40 coordinates representing the lineage-through-time (LTT) plot. To capture more information on the phylogenies generated by the BDSS model, we further enriched these SS with 14 new statistics on transmission chains describing the distribution of the duration between consecutive transmissions (internal tree nodes). Our SS are diverse, complementary and somewhat redundant. We used feed-forward neural networks (FFNN) with several hidden layers (**Fig. 2 b (i)**) that select and combine relevant information from the input features. In addition to SS, we provide both the tree size (*i.e.,* number of tips) and the sampling probability used to generate the tree, as input to our FFNN (**Fig. 2 a (vi)**). We will refer to this method as FFNN-SS.

**Fig. 2:**
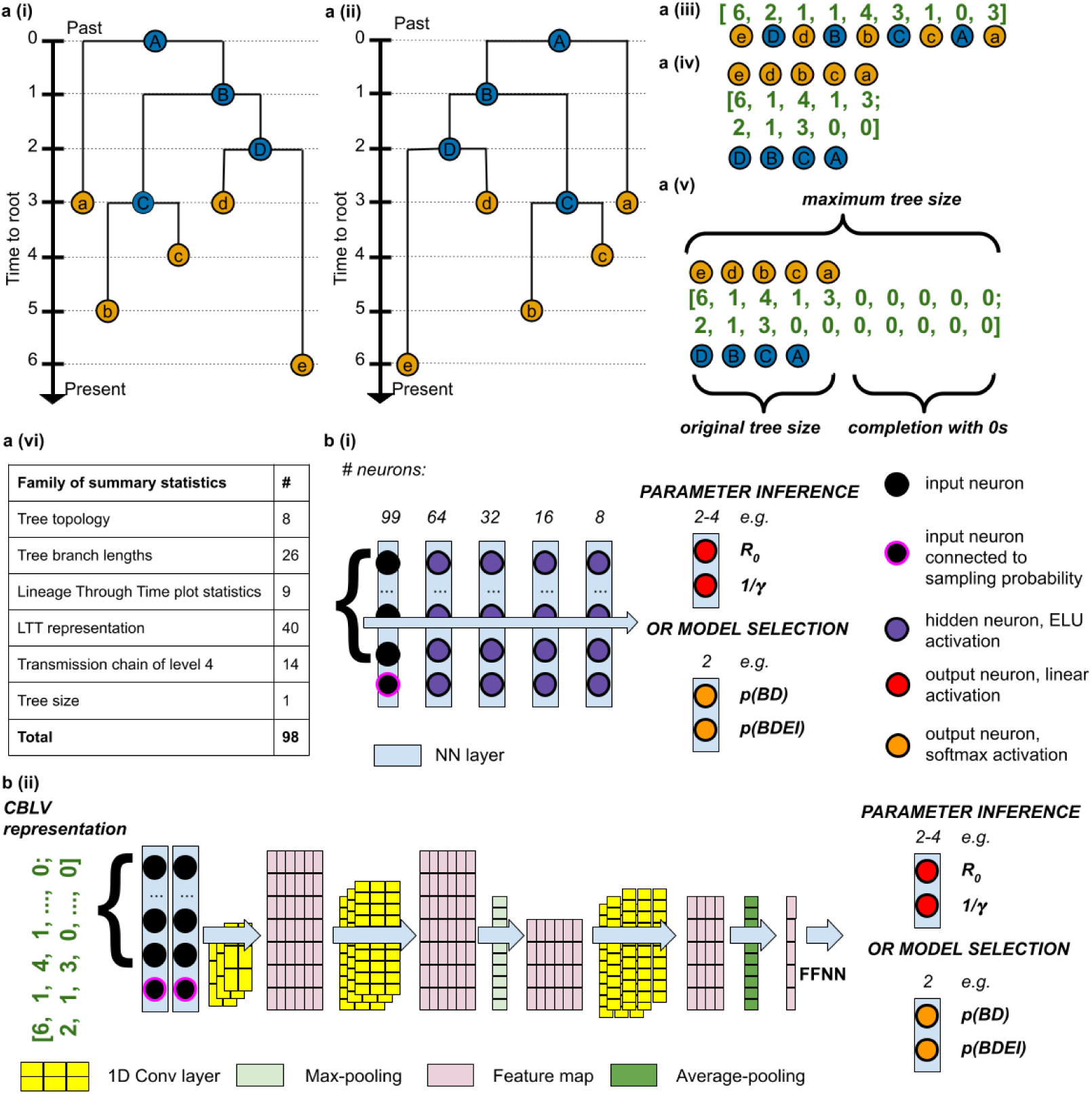
Pipeline for training neural networks on phylogenies. **Note to Fig.2. Tree representations**: **a (i),** simulated binary trees. Under each model from Fig. 1, we simulate many trees of variable size (50 to 200 tips for ‘small trees’ and 200 to 500 tips for ‘large trees’). For illustration, we have here a tree with 5 tips. We encode the simulations into two representations, either **a (ii-v)**, in a complete and compact tree representation called ‘Compact Bijective Ladderized Vector’ abbreviated as CBLV or a (vi) with summary statistics (SS). CBLV is obtained through **a (ii)** ladderization or sorting of internal nodes so that the branch supporting the most recent leaf is always on the left and **a (iii)** an inorder tree traversal, during which we append to a real-valued vector for each visited internal node its distance to the root and for each visited tip its distance to the previously visited internal node. We reshape this representation into **a (iv)**, an input matrix in which the information on internal nodes and leaves is separated into two rows. Finally, **a (v)**, we complete this matrix with zeros so that the matrices for all simulations have the size of largest simulation matrices. For illustration purpose, we here consider that the maximum tree size covered by simulations is 10, and the representation is thus completed with 0s accordingly. SS consists of **a (vi),** a set of 98 statistics: 83 published in *Saulnier et al*^[19]^, 14 on transmission chains and 1 on tree size. The information on sampling probability is added to both representations. **b: Neural networks** are trained on these representations to estimate parameter values or to select the underlying model. For SS, we use, **b (i)**, a deep feed-forward neural network (FFNN) of funnel shape (we show the number of neurons above each layer). For the CBLV representation we train, **b (ii),** Convolutional Neural Networks (CNN). The CNN is added on top of the FFNN. The CNN combines convolutional, maximum pooling and global average pooling layers, as described in detail in **Methods**.

### Compact vectorial tree representation

While converting raw information in the form of a phylogenetic tree into a set of SS, information loss is unavoidable. This means not only that the tree cannot be fully reconstructed from its SS, but also that depending on how much useful and relevant information is contained in the SS, the neural network may fail to solve the problem at hand. As an alternative strategy to SS, and to prevent information loss in the tree representation, we developed a representation called ‘Compact Bijective Ladderized Vector’ (CBLV).

Several vectorial representations of trees based either on polynomial^[26, 27]^, Laplacian spectrum^[28]^ or F matrices^[29]^ have been developed previously. However, they represent the tree shape but not the branch lengths^[26]^ or may lose information on trees^[28]^. In addition, some of these representations require vectors or matrices of quadratic size with respect to the number of tips^[29]^, or are based on complex coordinate systems of exponential size^[27]^.

Inspired by these approaches, we designed our concise, easily computable, compact, and bijective (i*.e.* 1-to-1) tree representation that applies to trees of variable size and is appropriate as machine learning input. To obtain this representation, we first ladderize the tree, that is, for each internal node, the descending subtree containing the most recently sampled tip is rotated to the left, **Fig. 2 a (ii)**. This ladderization step does not change the tree but facilitates learning by standardizing the input data. Moreover, it is consistent with trees observed in real epidemiological datasets, for example Influenza, where ladder-like trees reflect selection and are observed for several pathogens^[1]^. Then, we perform an inorder traversal^[30]^ of the ladderized tree, during which we collect in a vector for each visited internal node its distance to the root and for each tip its distance to the previously visited internal node. In particular, the first vector entry corresponds to the tree height. This transformation of a tree into a vector is bijective, in the sense that we can unambiguously reconstruct any given tree from its vector representation (**Supplementary Fig. 1**). The vector is as compact as possible, and its size grows linearly with the number of tips. We complete this vector with zeros to reach the representation length of the largest tree contained in our simulation set, and we add the sampling probability used to generate the tree (or an estimate of it when analysing real data; **Fig. 2 a (v), b (i))**.

Bijectivity combined with ladderization facilitates the training of neural networks, which do not need to learn that different representations correspond to the same tree. However, unlike our SS, this full representation does not have any high-level features. In CBLV identical subtrees will have the same representation in the vector whenever the roots of these subtrees have the same height, while the vector representation of the tips in such subtrees will be the same no matter the height of the subtree’s root. Similar subtrees will thus result in repeated patterns along the representation vector. We opted for Convolutional Neural Networks (CNN), which are designed to extract information on patterns in raw data. Our CNN architecture (**Fig. 2 b (ii)**) includes several convolutional layers that perform feature extraction, as well as maximum and average pooling layers that select relevant features and keep feature maps of reasonable dimensions. The output of the CNN is then fed into a FFNN that combines the patterns found in the input to perform predictions. In the rest of the manuscript, we refer to this method as CNN-CBLV.

### Simulated datasets

For each phylodynamic model (BD, BDEI, BDSS), we simulated 4 million trees, covering a large range of values for each parameter of epidemiological interest (R_0_, infectious period: 1/γ, incubation period: 1/ε, the fraction at equilibrium of superspreading individuals: f_SS_, and the superspreading transmission ratio: X_SS_). Of the 4 million trees, 3.99 million were used as a training set, and 10,000 as a validation set for early stopping in the training phase^[31]^. Additionally, we simulated another 10,000 trees, which we used as a testing set, out of which 100 were also evaluated with the gold standard methods, BEAST2 and TreePar, which are more time consuming. Another 1 million trees were used to define confidence intervals for estimated parameters. For BD and BDEI we considered two settings: one with small trees (50 to 199 tips, in **Supplementary Fig. 2**) and a second with large trees (200 to 500 tips, **Fig. 3**). For BDSS, we considered only the setting with large trees, as the superspreading individuals are at a low fraction and cannot be detected in small trees (results not shown). Lastly, we investigated the applicability of our approach to very large data sets, which are increasingly common with viral pathogens. To this goal, we generated for each model 10,000 ‘huge’ trees, with 5,000 to 10,000 tips each and with the same parameter ranges as used with the small and large trees. To estimate the parameter values of a huge tree, we extracted a nearly complete coverage of this tree by disjoint subtrees with 50 to 500 leaves. Then, we predicted the parameter values for every subtree using our NNs, and averaged subtree predictions to obtain parameter estimates for the huge tree.

**Fig. 3:**
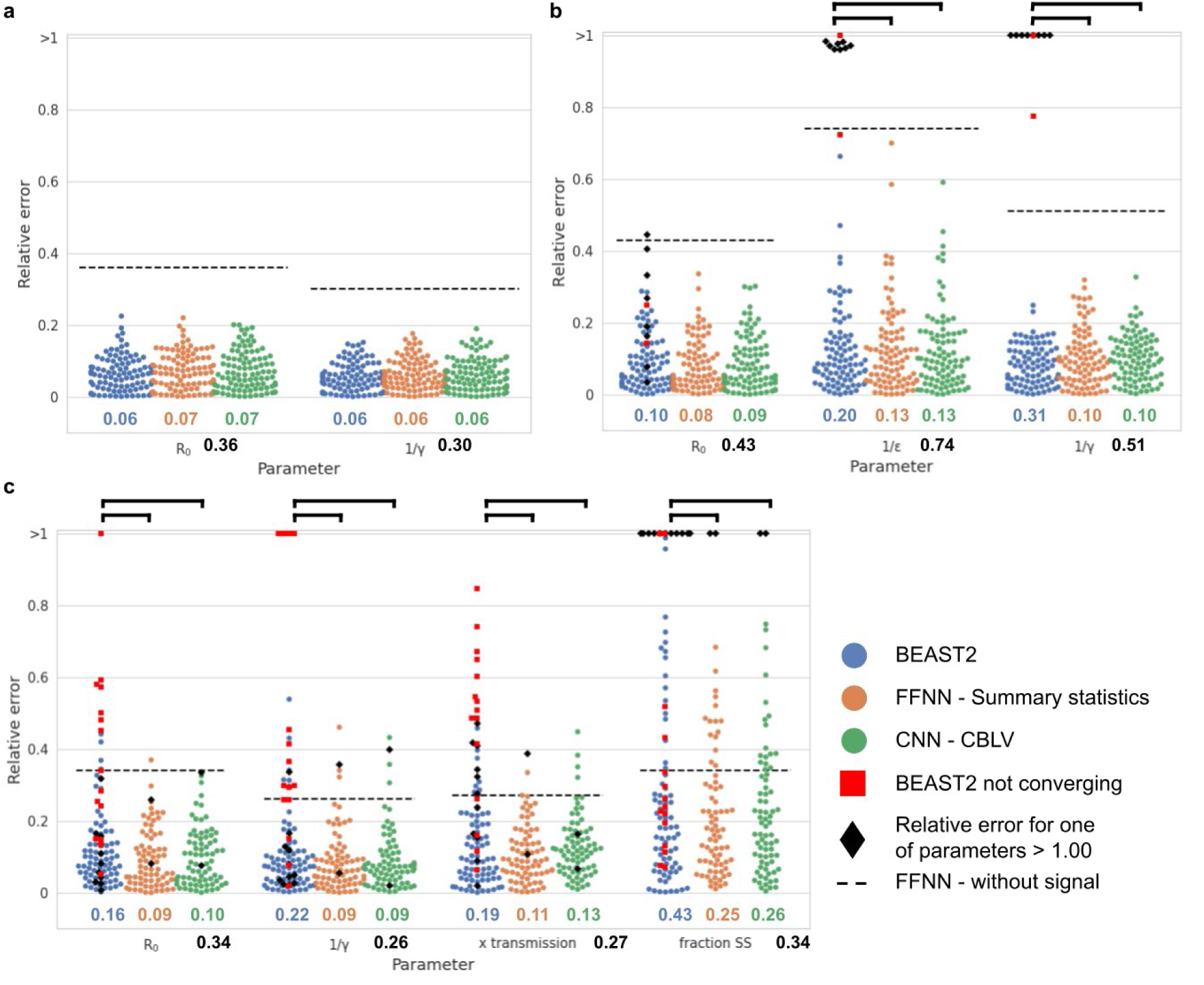
Assessment of deep learning accuracy. **Note to Fig. 3.** Comparison of inference accuracy by BEAST2 (in blue), deep neural network trained on SS (in orange) and convolutional neural network trained on the CBLV representation (in green) on 100 test trees. The size of training and testing trees was uniformly sampled between 200 and 500 tips. We show the relative error for each test tree. The error is measured as the normalized distance between the median *a posteriori* estimate by BEAST2 or point estimates by neural networks and the target value for each parameter. We highlight simulations for which BEAST2 did not converge and whose values were thus set to median of the parameter subspace used for simulations by depicting them as red squares. We further highlight the analyses with a high relative error (>1.00) for one of the estimates as black diamonds. We compare the relative errors for **a,** BD-simulated, **b,** BDEI-simulated and **c,** BDSS-simulated trees. Average relative error is displayed for each parameter and method in corresponding colour below each figure. The average error of a FFNN trained on summary statistics but with randomly permuted target is displayed as black dashed line and its value is shown in bold black below the x-axis. The accuracy of each method is compared by paired z-test; *P* < 0.05 is shown as thick full line; non-significant is not shown.

To increase the generality of our approach and avoid the arbitrary choice of the time scale (one unit can be a day, a week, or a year), we rescaled all trees and corresponding epidemiological parameters, such that the average branch length in a tree was equal to 1. After inference, we rescaled the estimated parameter values back to the original time scale.

### Neural networks yield more accurate parameter estimates than gold standard methods

We compared accuracy of parameter estimates yielded by our deep learning methods and those yielded by two state-of-the-art phylodynamics inference tools, BEAST2^[12, 13]^ and TreePar^[5]^. The comparison shows that our deep learning methods trained with SS and CBLV are either comparable (BD) or more accurate (BDEI and BDSS) than the state-of-the-art inference methods (**Fig. 3, Supplementary Tab. 1**). The simple BD model has a closed form solution for the likelihood function, and thus BEAST2 results are optimal in theory^[8, 9]^. Our results with BD are similar to those obtained with BEAST2, and thus nearly optimal as well. For BDEI and BDSS our results are more accurate than BEAST2, which is likely explained by numerical approximations of likelihood calculations in BEAST2^[5,10,11]^ for these models. These approximations can lead BEAST2 to a lack of convergence (2% cases for BDEI and 15% cases for BDSS) or a convergence to local optima. We suspect BEAST2 of converging to local optima when it converged to values with high relative error (>1.0; 8% cases for BDEI and 11% cases for BDSS, **Fig. 3 b-c**). Furthermore, our deep learning approaches showed a lower bias in parameter estimation than BEAST2 (**Supplementary Tab. 2**). As expected, both approaches, FFNN-SS and CNN-CBLV, get more accurate with larger trees (**Supplementary Fig. 3**).

We tried to perform maximum likelihood estimation (MLE) implemented in the TreePar package^[5]^ on the same trees as well. While MLE under BD model on simulations yielded as accurate results as BEAST2, for more complex models it showed overflow and underflow issues (*i.e.,* reaching infinite values of likelihood) and yielded inaccurate results, such as more complex models (BDEI, BDSS) having lower likelihood than a simpler, nested one (BD) for a part of simulations (results not shown). These issues were more prominent for larger trees. TreePar developers confirmed these limitations and suggested using the latest version of BEAST2 instead.

To further explain the performance of our NNs, we computed the likelihood value of their parameter estimates. This was easy with the BD model since we have a closed form solution for the likelihood function. The results with this model (**Supplementary Tab. 3**, using TreePar) showed that the likelihoods of both FFNN-SS and CNN-CBLV estimates are similar to BEAST2’s, which explains the similar accuracy of the three methods (**Fig. 3**). We also computed the likelihood of the ‘true’ parameter values used to simulate the trees, in order to have an independent and solid assessment. If a given method tends to produce higher likelihood than that of the true parameter values, then it performs well in terms of likelihood optimization, as optimizing further should not result in higher accuracy. The results (**Supplementary Tab. 3**) were again quite positive, as BEAST2 and our NNs achieved a higher likelihood than the true parameter values for ∼70% of the trees, with a significant mean difference. With BDEI and BDSS, applying the same approach proved difficult due to convergence and numerical issues, with both BEAST2 and TreePar (see above). For the partial results we obtained (not shown), the overall pattern seems to be similar to that with BD: the NNs obtain highly likely solutions, with similar likelihood as BEAST2’s (when it converges and produces reasonable estimates), and significantly higher likelihood than that of the true parameter values. All these results are remarkable, as the NNs do not explicitly optimize the likelihood function associated to the models, but use a radically different learning approach, based on simulation.

### Neural networks are fast inference methods

We compared the computing time required by each of our inference methods. All computing times were estimated for a single thread of our cluster, except for the training of neural architectures where we used our GPU farm. Neural networks require heavy computing time in the learning phase; for example, with BDSS (the most complex model), simulating 4M large trees requires ∼800 CPU hours, while training FFNN-SS and CNN-CBLV requires ∼5 and ∼150 hours, respectively. However, with NNs, inference is almost instantaneous and takes ∼0.2 CPU seconds per tree on average, including encoding the tree in SS or CBLV, which is the longest part. For comparison, BEAST2 inference under the BD model with 5 million MCMC steps takes on average ∼0.2 CPU hours per tree, while inference under BDEI and BDSS with 10 million MCMC steps takes ∼55 CPU hours and ∼80 CPU hours per tree, respectively. In fact, the convergence time of BEAST2 is usually faster (∼6 CPU hours with BDEI and BDSS), but can be very long in some cases, to the point that convergence is not observed after 10 million steps (see above).

### Neural networks have high generalization capabilities and apply to very large data sets

In statistical learning theory^[31]^, generalization relates to the ability to predict new samples drawn from the same distribution as the training instances. Generalization is opposed to rote learning and overfitting, where the learned classifier or regressor predicts the training instances accurately, but new instances extracted from the same distribution or population poorly. The generalization capabilities of our NNs were demonstrated, as we used independent testing sets in all our experiments (**Fig. 3**). However, we expect poor results with trees that depart from the training distribution, for example showing very high R_0_, while our NNs have been trained with R_0_ in the range ^[1, 5]^. If, for a new study, larger or different parameter ranges are required, we must retrain the NNs with *ad hoc* simulated trees. However, a strength of NNs is that thanks to their flexibility and approximation power, very large parameter ranges can be envisaged, to avoid repeating training sessions too often.

Another sensible issue is that of the size of the trees. Our NNs have been trained with trees of 50-to-199 tips (small) and 200-to-500 tips (large), that is, trees of moderate size (but already highly time consuming in a Bayesian setting, for the largest ones). Thus, we tested the ability to predict the parameters of small trees using NNs trained on large trees, and vice versa, the ability to predict large trees with NNs trained on small trees. The results (**Supplementary Fig. 4**) are surprisingly good, especially with summary statistics (FFNN-SS) which are little impacted by these changes of scale as they largely rely on means (*e.g.*, of branch lengths^[19]^). This shows unexpected generalization capabilities of the approach regarding tree size. Most importantly, the approach can accurately predict huge trees (**Fig. 4**) using their subtrees and the means of the corresponding parameter estimates, in ∼1 CPU minute. This extends the applicability of the approach to data sets that cannot be analysed today, unless using similar tree decomposition and very long calculations to analyse all subtrees.

**Fig. 4:**
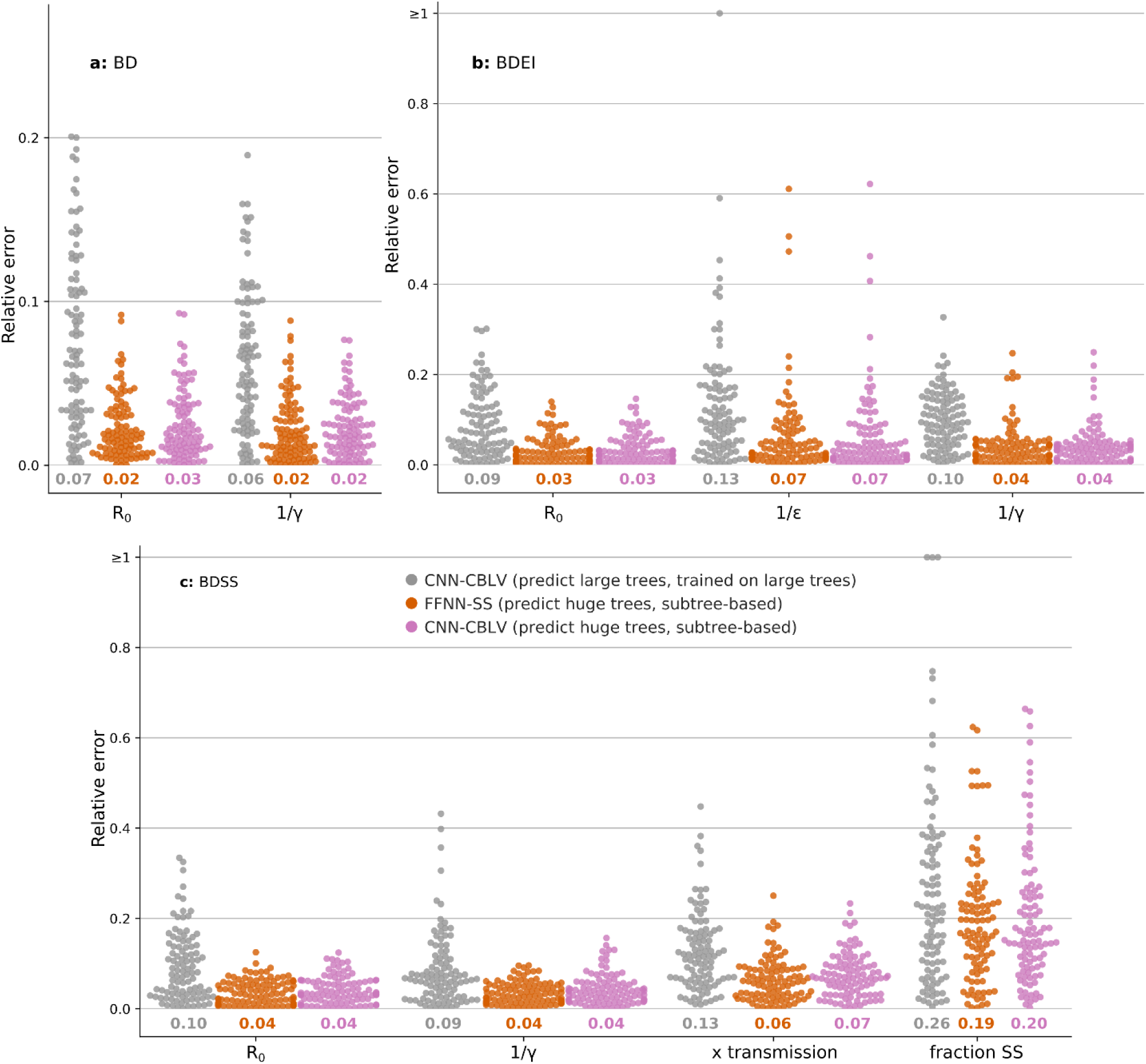
Deep learning accuracy with ‘huge’ trees. **Note to Fig. 4.** Comparison of inference accuracy by neural networks trained on large trees in predicting large trees (CNN-CBLV, in grey, same as in Fig. 3) and huge trees (FFNN-SS, in orange, and CBLV-NN, in pink) on 100 large and 100 huge test trees. The training and testing large trees are the same as in Fig. 3 (between 200 and 500 tips each). The huge testing trees were generated for the same parameters as the large training and testing trees, but their size varied between 5,000 and 10,000 tips. We show the relative error for each test tree. The error is measured as the normalized distance between the point estimates by neural networks and the target values for each parameter. We compare the relative errors for **a**, BD-simulated, **b**, BDEI-simulated and **c**, BDSS-simulated trees. Average relative error is displayed for each parameter and method in corresponding colour below each plot.

### Neural networks are accurate methods for model selection

We trained CNN-CBLV and FFNN-SS on simulated trees to predict the birth-death model under which they were simulated (BD or BDEI for small trees; BD, BDEI or BDSS for large trees). Note that for parameters shared between multiple models, we used identical parameter value ranges across all these models (**Supplementary Tab. 4**). Then, we assessed the accuracy of both of our approaches on 100 simulations obtained with each model and compared it with the model selection under BEAST2 based on Akaike information criterion through Markov Chain Monte Carlo (AICM)^[32, 33]^. The AICM, similar to deviance information criterion (DIC) by Gelman *et al.*^[32]^, does not add computational load and is based on the average and variance of posterior log-likelihoods along the Markov Chain Monte Carlo (MCMC).

FFNN-SS and CNN-CBLV have similar accuracy (**Supplementary Tab. 5**), namely 92% for large trees (BD vs BDEI vs BDSS), and accuracy of 91% and 90%, respectively, for small trees (BD vs BDEI). BEAST2 yielded an accuracy of 91% for large trees and 87% for small trees. The non-converging simulations were not considered for any of these methods (*i.e.,* 5% simulations for small trees and 24% for large trees).

The process of model selection with a neural network is as fast as the parameter inference (∼0.2 CPU seconds per tree). This represents a practical, fast and accurate way to perform model selection in phylodynamics.

### Neural networks are well suited to learn complex models

To assess the complexity of learned models, we explored other inference methods, namely: (1) linear regression as a baseline model trained on summary statistics (LR-SS); (2) FFNN trained directly on CBLV (FFNN-CBLV); (3) CNN trained on Compact Random Vector (CNN-CRV), for which the trees were randomly rotated, instead of being ladderized as in **Fig. 2 (ii)**; and (4) two “null models”.

LR-SS yielded inaccurate results even for the BD model (**Supplementary Tab. 1**), which seems to contrast with previous findings^[19]^, where LR approach combined with ABC performed only slightly worse than BEAST2. This can be explained by the lack of rejection step in LR-SS, which enables to locally reduce the complexity of the relation between the representation and the inferred values to a linear one^[18]^. However, the rejection step requires a metric (*e.g.,* the Euclidean distance), which may or may not be appropriate depending on the model and the summary statistics. Moreover, rejection has a high computational cost with large simulation sets.

Neural networks circumvent these problems with rejection and allow for more complex, non-linear relationships between the tree representation and the inferred values to be captured. This is also reflected in our results with FFNN-CBLV and CNN-CRV, which both proved to be generally more accurate than LR-SS. However, FFNN-CBLV was substantially less accurate than CNN-CBLV (**Supplementary Tab. 1**, **Supplementary Fig. 5**). This indicates the presence of repeated patterns that may appear all along the vectorial representation of trees, such as subtrees of any size, which are better extracted by CNN than by FFNN. In its turn, CNN-CRV required larger training sets to reach an accuracy comparable to CNN-CBLV (**Supplementary Tab. 1**, **Supplementary Fig. 5**), showing that the ladderization and bijectivity of the CBLV helped the training.

To assess how much information is actually learned, we also measured the accuracy of two “null models”: FFNN trained to predict randomly permuted target values; and a random predictor, where parameter values were sampled from prior distributions. Results show that the neural networks extract a considerable amount of information for most of the estimated parameters (**Supplementary Tab. 1**). The most difficult parameter to estimate was the fraction of superspreading individuals in BDSS model, with accuracy close to random predictions with small trees, but better performance as the tree size increases (**Fig. 4, Supplementary Fig. 3**).

### SS is simpler, but CBLV has high potential for application to new models

FFNN-SS and CNN-CBLV show similar accuracy across all settings (**Fig. 3, Supplementary Tab. 1-2**), including when predicting huge trees from their subtrees (**Fig. 4**). The only exception is the prediction of large trees using NNs trained with small trees (**Supplementary Fig. 4**), where FFNN-SS is superior to CNN-CBLV, but this goes beyond the recommended use of the approach, as only a part of the (large) query tree is given to the (small) CNN-CBLV.

However, the use of the two representations is clearly different, and it is likely that with new models and scenarios their accuracy will differ. SS requires a simpler architecture (FFNN) and is trained faster (*e.g.*, 5 hours with large BDSS trees), with less training instances (**Supplementary Fig. 6**). However, this simplicity is obtained at cost of a long preliminary work to design appropriate summary statistics for each new model, as was confirmed in our analyses of BDSS simulations. To estimate the parameters of this model, we added summary statistics on transmission chains on top of the SS taken from Saulnier *et al.*^[19]^. This improved the accuracy of superspreading fraction estimates of the FFNN-SS, so that it was comparable to the CNN-CBLV, while the accuracy for the other parameters remained similar (**Supplementary Fig. 7**). The advantage of the CBLV is its generality, meaning there is no loss of information between the tree and its representation in CBLV regardless of which model the tree was generated under. However, CBLV requires more complex architectures (CNN), more computing time in the learning phase (150 hours with large BDSS trees) and more training instances (**Supplementary Fig. 6**). Such an outcome is expected. With raw CBLV representation, the convolutional architecture is used to “discover” relevant summary statistics (or features, in machine learning terminology), which has a computational cost.

In fact, the two representations should not be opposed. An interesting direction for further research would be to combine them (*e.g.* during the FFNN phase), to possibly obtain even better results. Moreover, SS are still informative and useful (and quickly computed), in particular to perform sanity checks, both a priori and a posteriori (**Fig. 5, Supplementary Fig. 8**), or to quickly evaluate the predictability of new models and scenarios.

**Fig. 5:**
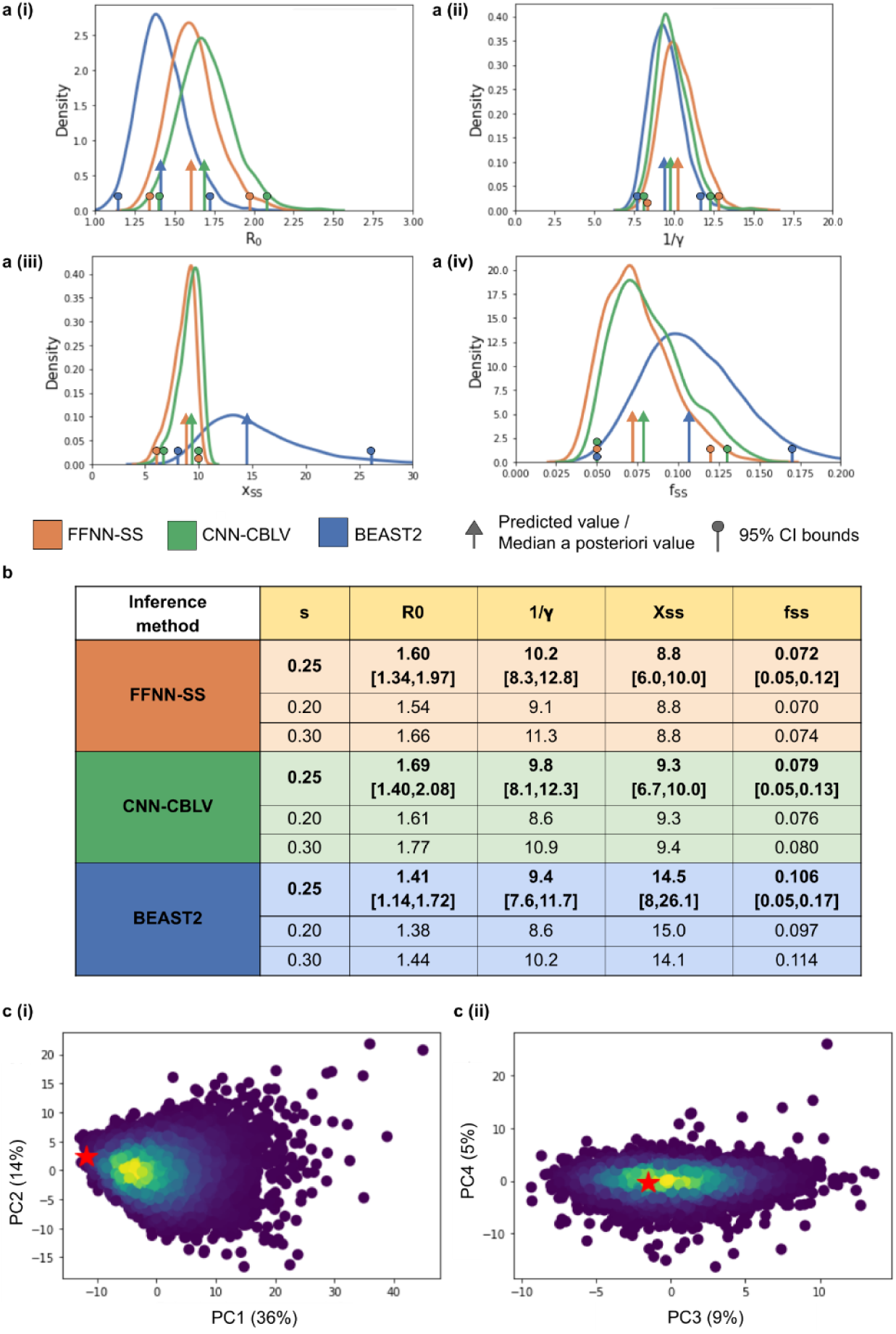
Parameter inference on HIV data sampled from MSM Zurich. **Note to Fig. 5.** Using BDSS model with BEAST2 (in blue), FFNN-SS (in orange), and CNN-CBLV (in green) we infer, **a (i),** basic reproduction number, **a (ii),** infectious period (in years), **a (iii),** superspreading transmission ratio and, **a (iv),** superspreading fraction. For FFNN-SS and CNN-CBLV, we show the posterior distributions and the 95% CIs obtained with a fast approximation of the parametric bootstrap (**Methods**). For BEAST2, the posterior distributions and 95% CI were obtained considering all reported steps (9,000 in total) excluding the 10% burn-in. Arrows show the position of the original point estimates obtained with FFNN-SS and CNN-CBLV and the median *a posteriori* estimate obtained with BEAST2. Circles show lower and upper boundaries of 95% CI. **b,** these values are reported in a table, together with point estimates obtained while considering lower and higher sampling probabilities (0.20 and 0.30). **c,** 95% CI boundaries obtained with FFNN-SS are used to perform an *a posteriori* model adequacy check. We simulated 10,000 trees with BDSS while resampling each parameter from a uniform distribution, whose upper and lower bounds were defined by the 95% CI. We then encoded these trees into SS, performed PCA and projected SS obtained from the HIV MSM phylogeny (red stars) on these PCA plots. We show here the projection into **c (i),** first two components of PCA, **c (ii),** the 3^rd^ and 4^th^ components, together with the associated percentage of variance displayed in parentheses. Warm colours correspond to high density of simulations.

### Showcase study of HIV in MSM subpopulation in Zurich

The Swiss HIV Cohort is densely sampled, including more than 16,000 infected individuals^[24]^. Datasets extracted from this cohort have often been studied in phylodynamics^[8, 25]^. We analysed a dataset of an MSM subpopulation from Zurich, which corresponds to a cluster of 200 sequences studied previously by Rasmussen *et al.*^[25]^, who focused on the degree of connectivity and its impact on transmission between infected individuals. Using coalescent approaches, they detected the presence of highly connected individuals at the beginning of the epidemic and estimated R_0_ to be between 1.0 and 2.5. We used their tree as input for neural networks and BEAST2.

To perform analyses, one needs an estimate of the sampling probability. We considered that: (1) the cohort is expected to include around 45% of Swiss individuals infected with HIV^[24]^; and (2) the sequences were collected from around 56% of individuals enrolled in this cohort^[34]^. We used these percentages to obtain an approximation of sampling probability of 0.45*0.56 ∼ 0.25 and used this value to analyse the MSM cluster. To check the robustness of our estimates, we also used sampling probabilities of 0.2 and 0.3 in our estimation procedures.

First, we performed a quick sanity check considering the resemblance of HIV phylogeny with simulations obtained with each model. Two approaches were used, both based on SS (**Supplementary Fig. 8**). Using principal component analysis (PCA), all three considered birth-death models passed the check. However, when looking at the 97 SS values in detail, namely checking whether the observed tree SS were within the [min, max] range of the corresponding simulated values, the BD and BDEI models were rejected for some of the SS (5 for both models, all related to branch lengths). Then, we performed model selection (BD vs BDEI vs BDSS) and parameter estimation using our two methods and BEAST2 (**Fig. 5 a-b**). Finally, we checked the model adequacy with a second sanity check, derived from the inferred values and SS (**Fig. 5 c, Supplementary Fig. 8)**.

Model selection with CNN-CBLV and FFNN-SS resulted in the acceptance of BDSS (probability of 1.00 versus 0.00 for BD and BDEI), and the same result was obtained with BEAST2 and AICM. These results are consistent with our detailed sanity check, and with what is known about HIV epidemiology, namely, the presence of superspreading individuals in the infected subpopulation^[35]^ and the absence of incubation period without infectiousness such as is emulated in BDEI^[36]^.

We then inferred parameter values under the selected BDSS model (**Fig. 5 a-b**). The values obtained with FFNN-SS and CNN-CBLV are close to each other, and the 95% CI are nearly identical. We inferred an R_0_ of 1.6 and 1.7, and an infectious period of 10.2 and 9.8 years, with FFNN-SS and CNN-CBLV, respectively. Transmission by superspreading individuals was estimated to be around 9 times higher than by normal spreaders and superspreading individuals were estimated to account for around 7-8% of the population. Our R_0_ estimates are consistent with the results of a previous study^[8]^ performed on data from the Swiss cohort, and the results of Rasmussen *et al.*^[25]^ with this dataset. The infectious period we inferred is a bit longer than that reported by *Stadler et al*, who estimated it to be 7.74 [95% CI 4.39-10.99] years^[8]^. The infectious period is a multifactorial parameter depending on treatment efficacy and adherence, the times from infection to detection and to the start of treatment, *etc.* In contrast to the study by *Stadler et al*, whose data were sampled in the period between 1998 and 2008, our dataset covers also the period between 2008 and 2014, during which life expectancy of patients with HIV was further extended^[37]^. This may explain why we found a longer infectious period (with compatible CIs). Lastly, our findings regarding superspreading are in accordance with those of Rassmussen *et al.*^[25]^, and with a similar study in Latvia^[5]^ based on 40 MSM sequences analysed using a likelihood approach. Although the results of the latter study may not be very accurate due to the small dataset size, they still agree with ours, giving an estimate of a superspreading transmission ratio of 9, and 5.6% of superspreading individuals. Our estimates were quite robust to the choice of sampling probability (*e.g.,* R0 = 1.54, 1.60 and 1.66, with FFNN-SS and a sampling probability of 0.20, 0.25 and 0.30, respectively, **Fig. 5 b**).

Compared to BEAST2, the estimates of the infectious period and R0 were similar for both approaches, but BEAST2 estimates were higher for the transmission ratio (14.5) and the superspreading fraction (10.6%). These values are in accordance with the positive bias of BEAST2 estimates that we observed in our simulation study for these two parameters, while our estimates were nearly unbiased (**Supplementary Tab. 2**).

Finally, we checked the adequacy of BDSS model by resemblance of HIV phylogeny to simulations. Using inferred 95% CI, we simulated 10,000 trees and performed PCA on SS, to which we projected the SS of our HIV phylogeny. This was close to simulations, specifically close to the densest swarm of simulations, supporting the adequacy of both the inferred values and the selected model (**Fig. 5 c**). When looking at the 97 SS in detail, some of the observed values where not in the [min, max] range of the 10,000 simulated values. However, these discordant SS were all related to the lineage-through-time plot (LTT; *e.g.*, *x* and *y* coordinates of this plot; **Supplementary Fig. 8**), consistent with the fact that the probabilistic, sampling component of the BDSS model is an oversimplification of actual sampling schemes, which depend on contact tracing, sampling campaigns and policies, etc.

## DISCUSSION AND PERSPECTIVES

In this manuscript, we presented new methods for parameter inference and model selection in phylodynamics based on deep learning from phylogenies. Through extensive simulations, we established that these methods are at least as accurate as state-of-the-art methods and capable of predicting very large trees in minutes, which cannot be achieved today by any other existing method. We also applied our deep learning methods to the Swiss HIV dataset from MSM and obtained results consistent with current knowledge of HIV epidemiology.

Using BEAST2, we obtained inaccurate results for some of the BDEI and BDSS simulations. While BEAST2 has been successfully deployed on many models and tasks, it clearly suffers from approximations in likelihood computation with these two models. However, these will likely improve in near future. In fact, we already witnessed substantial improvements done by BEAST2 developers to the BDSS model, while carrying out this research.

Both of our neural network approaches circumvent likelihood computation and thereby represent a new way of using molecular data in epidemiology, without the need to solve large systems of differential equations. This opens the door to novel phylodynamics models, which would make it possible to answer questions previously too complex to ask. This is especially true for CBLV representation, which does not require the design of new summary statistics, when applied to trees generated by new mathematical models. A direction for further research would be to explore such models, for example based on structured coalescent^[38, 39]^, or to extend the approach to macroevolution and species diversification models^[40]^, which are closely related to epidemiological models. Other fields related to phylodynamics, such as population genetics, have been developing likelihood-free methods^[41]^, for which our approach might serve as a source of inspiration.

A key issue in both phylodynamics and machine learning applications is scalability. Our results show that very large phylogenies can be analysed very efficiently (∼1 minute for 10,000 tips), with resulting estimates more accurate than with smaller trees (**Fig. 4**), as predicted by learning theory. Again, as expected, more complex models require more training instances, especially BDSS using CBLV (**Supplementary Fig. 3**), but the ratio remains reasonable, and it is likely that complex (but identifiable) models will be handled efficiently with manageable training sets. Surprisingly, we did not observe a substantial drop of accuracy with lower sampling probabilities (results not shown). To analyse very large trees, we used a decomposition into smaller, disjoint subtrees. In fact, all our NNs were trained with trees of moderate size (<500 tips). Another approach would be to learn directly from large trees. This is an interesting direction for further research, but this poses several difficulties. The first is that we need to simulate these very large trees, and a large number of them (millions or more). Then, SS is the easiest representation to learn, but at the risk of losing essential information, which means that new summary statistics will likely be needed for sufficiently complete representation of very large phylogenies. Similarly, with CBLV more complex NN architectures (*e.g.*, with additional and larger kernels in the convolutional layers) will likely be needed, imposing larger training sets. Combining both representations (*e.g.*, during the FFNN phase) is certainly an interesting direction for further research. Note, however, that the predictions of both approaches for the three models we studied are highly correlated (Pearson coefficient nearly equal to 1 for most parameters), which means that there is likely little room for improvement (at least with these models).

A key advantage of the deep learning approaches is that they yield close to immediate estimates and apply to trees of varying size. Collection of pathogen genetic data became standard in many countries, resulting in densely sampled infected populations. Examples of such datasets include HIV in Switzerland and UK^[24, 42]^, 2013 Ebola epidemics^[6]^, several Influenza epidemics and the 2019 SARS-Cov-2 pandemic (www.gisaid.org)^[43]^. For many such pathogens, trees can be efficiently and accurately inferred^[44–46]^ and dated^[47–49]^ using standard approaches. When applied to such dated trees, our methods can perform model selection and provide accurate phylodynamic parameter estimates within a fraction of a second. Such properties are desirable for phylogeny-based real-time outbreak surveillance methods, which must be able to cope with the daily influx of new samples, and thus increasing size of phylogenies, as the epidemic unfolds, in order to study local outbreaks and clusters, and assess and compare the efficiency of healthcare policies deployed in parallel. Moreover, thanks to the subtree picking and averaging strategy, it is now possible to analyse extremely large phylogenies, and the approach could be used to track the evolution of parameters (*e.g.*, R_0_) in different regions (sub-trees) of a global tree, as a function of dates (as in Bayesian skyline models^[4]^), geographical areas, viral variants etc.

## Supporting information

Voznica_et_al_Supplementary_information

## ACKNOWLEDGEMENT

We would like to thank Dr Kary Ocaña and Tristan Dot for initiating experiments on machine learning and phylogenetic trees in our laboratory. We would like to thank Quang Tru Huynh for administrating the GPU farm at Institut Pasteur and the INCEPTION program (Investissement d’Avenir grant ANR-16-CONV-0005) that financed the GPU farm. We would like to thank Dr Christophe Zimmer from Institut Pasteur, Sophia Lambert and Dr Hélène Morlon from Institut de Biologie de l’Ecole Normale Supérieure IBENS and Dr Guy Baele from Katholieke Universiteit KU Leuven for useful discussions and Dr Isaac Overcast from IBENS for critical reading of the manuscript. We would like to thank Dr Tanja Stadler and Jérémie Sciré for their help with BEAST2 and MLE approaches. JV is supported by Ecole Normale Supérieure Paris-Saclay and by ED Frontières de l’Innovation en Recherche et Education, Programme Bettencourt. VB would like to thank Swiss National Science Foundation for funding (Early PostDoc mobility grant P2EZP3_184543). OG is supported by PRAIRIE (ANR-19-P3IA-0001).

## METHODS

### TREE REPRESENTATION USING SUMMARY STATISTICS (SS)

We use a set of 98 summary statistics (SS), to which we add the sampling probability, summing to a vector of 99 values.

#### Saulnier et al summary statistics

We use the 83 SS proposed by Saulnier *et al.*^[19]^:

- 8 SS on tree topology
- 26 SS on branch lengths
- 9 SS on Lineage-Through-Time (LTT) plot
- 40 SS providing the coordinates of the LTT plot

The computing time of these statistics grows linearly with tree size. For details, see the original paper.

#### Additional summary statistics

In addition to Saulnier *et al.*^[19]^ statistics, we designed 14 SS on transmission chains. Moreover, we provide the number of tips in the tree as input resulting in 83+14+1 = 98 SS in total.

The statistics on transmission chains are designed to capture information on the superspreading population. A superspreading individual transmits to more individuals within a given time period than a normal spreader. We thus expect that with superspreading individuals we would have shorter transmission chains. To have a proxy for the transmission chain length, we look at the sum of 4 subsequent shortest times of transmission for each internal node. This gives us a distribution of time-durations of 4-transmission chains. We assume that information on the transmission dynamics of superspreading individuals is retained in the lower (*i.e.*, left) tail of 4-transmission-chain lengths distribution, which contains relatively many transmissions with short time to next transmission, while the information on normal spreaders should be present in the rest of the distribution.

The implementation of these 4-transmission-chain SS is the following. For each internal node, we sum the distances from the internal node to its closest descendant nodes, descending exactly four times, that is, we take first the distance from the given internal node to its closest child node (of level 1), then from the (level 1) child node, we take its distance to its own closest child node (of level 2), *etc.* If one of the closest descendant nodes is a tip (except for the last one in the chain), we do not retain any value for the given internal node. Other options, like the shortest 4-edge pathway, could have been used as well and would likely give comparable results.

On the obtained distribution of 4-transmission-chain lengths, we compute 14 statistics:

- number of 4-transmission chains in the tree
- 9 deciles of 4-transmission-chain lengths distribution
- minimum and maximum values of 4-transmission-chain lengths distribution
- mean value of 4-transmission-chain lengths
- variance of 4-transmission-chain lengths

Adding the same summary statistics but on chains comprising 2, 3 and 5 consecutive transmissions had a negligible impact on parameter inference accuracy (data not shown).

### COMPLETE AND COMPACT TREE REPRESENTATION (CBLV)

Simulated dated trees are encoded in the form of real-valued vectors, which are then used as input for the neural networks. The representation of a tree with *n* tips is a vector of length 2*n*-1, where one single real-valued scalar corresponds to one internal node or tip. This representation thus scales linearly with the tree size. The encoding is achieved in two steps: tree ladderization and tree traversal.

#### Tree ladderization

The tree ladderization consists in ordering the children of each node. Child nodes are sorted based on the sampling time of the most recently sampled tip in their subtrees: for each node, the branch supporting the most recently sampled subtree is rotated to the left, as in **Fig. 2 a (i-ii)**.

We considered several alternatives with different criteria for child (subtree) sorting instead of ladderization: sampling time of the most anciently sampled tip, subtree length (*i.e.*, sum of all branch lengths including the rooting branch), diversification (*i.e.*, number of tips), normalized branch lengths (*i.e.*, subtree length divided by the number of tips), *etc*. These did not yield better results than CBLV. We show in **Supplementary Fig. 5** the comparison of CBLV with Compact Random Vector (CRV), for which internal nodes were sorted randomly before the tree traversal, showing that CRV yields poorer results than CBLV, as expected.

### Tree traversal and encoding

Once the tree is sorted, we perform an inorder tree traversal, using a standard recursive algorithm from the depth first family^[30]^. When visiting a tip, we add its distance to the previously visited internal node or its distance to the root, for the tip that is visited first (*i.e.*, the tree height due to ladderization). When visiting an internal node, we add its distance to the root. Examples of encoding are shown in **Fig. 2 a (ii-iii)**. This gives us the Compact Bijective Ladderized Vector (CBLV). We then separate information relative to tips and to internal nodes into two rows (**Fig. 2 a (iv)**) and complete the representation with zeros until reaching the size of the largest simulated tree for the given simulation set (**Fig. 2 a (v)**).

#### Properties of CBLV

CBLV has favourable features for deep learning. Ladderization does not actually change the input tree (phylogenies are unordered trees), but by ordering the subtrees it standardizes the input data and facilitates the learning phase, as observed with CRV (**Supplementary Fig. 5**). Then, the inorder tree traversal procedure is a bijective transformation, as it transforms a tree into a tree-compatible vector, from which the (ordered) tree can be reconstructed unambiguously, using a simple path-agglomeration algorithm shown in **Supplementary Fig. 1**. Note, however, that not all vectors are tree-compatible (*e.g.,* all entries must be non-negative; the second entry must be less than the first one, *etc*.). CBLV is “as concise as possible” being composed of 2*n*-1 real values (**Fig. 2 a (iii))**, where *n* is the number of tips. A rooted tree has 2*n*-2 branches, and thus 2*n*-2 entries are needed to represent the branch lengths. In our 2*n*-1 vectorial encoding of trees, we not only represent the branch lengths, but also the tree topology using only 1 additional entry.

The compactness and bijectivity of tree representation reduce the number of simulations required for training the neural network (**Supplementary Fig. 5**). This is because the number of parameters to be trained remains reasonable with compact representation. Moreover, the networks do not need to learn that several different inputs correspond to the same tree.

Our neural networks are intended to apply to trees of variable sizes (*e.g.*, trees of 200 to 500 tips in our experiments with ‘large’ trees). Thus, they are trained on representations of different lengths (*e.g.*, a vector of length 399 for a tree of 200 tips), that we complete with zeroes to reach the length of the largest trees (*i.e.*, 999 for 500 tips). We add an additional zero to obtain a two-row matrix (500*2 for 500 tips).

#### Alternative tree representations

Our CBLV tree representation could likely be improved to ease the learning phase and obtain even better parameter estimates. We tested several alternative representations, some inspired by the polynomial representation of small subtrees^[26, 27]^, the Laplacian spectrum^[28]^ and additive distance matrices that are equivalent to trees^[50]^. None was by far as convincing as CBLV, which is likely due to their large size (*e.g.*, *n*^2^ for distance matrices) or numerical instabilities and potential loss of information (*e.g.*, for Laplacian spectrum). Moreover, the margin for improvement of the accuracy of CNN-CBLV for the BD model, and likely for other models, is low. This is due to the observation that the accuracy of CNN-CBLV is similar to that of likelihood-based approaches for the BD model, and the fact that we have an analytical likelihood formula for the BD model, making the likelihood-based approach itself optimal^[8, 9]^.

### TREE RESCALING

Before encoding, the trees are rescaled so that the average branch length is 1, that is, each branch length is divided by the average branch length of the given tree, called rescale factor. The values of the corresponding time-dependent parameters (*i.e.*, infectious period and incubation period) are divided by the rescale factor too. The NN is then trained to predict these rescaled values. After parameter prediction, the predicted parameter values are multiplied by the rescale factor and thus rescaled back to the original time scale.

This step enables us to overcome problems of arbitrary time scales of input trees and makes a pre-trained NN more generally applicable. More specifically, an input tree with a time scale in days will be associated naturally with the same output as the same tree with a time scale in years, since both these trees will be rescaled to the same intermediate tree with average branch length of 1. Rescaling thus makes it possible to apply the same pre-trained NN to phylogenies reconstructed from sequences of a pathogen associated with an infectious period on the scale of days (*e.g.*, Ebola virus) or years (*e.g.*, HIV).

### REDUCTION AND CENTERING OF SUMMARY STATISTICS REPRESENTATION

Before training our NN and after having rescaled the trees to unit average branch length (see the sub-section above), we reduce and center every summary statistic by subtracting the mean and scaling to unit variance. To achieve this, we use the standard scaler from the scikit-learn package^[51]^, which is fitted to the training set. This does not apply to

CBLV representation, to avoid losing the ability to reconstruct the tree.

### PARAMETER AND MODEL INFERENCE USING NEURAL NETWORKS

We implemented deep learning methods in Python 3.6 using Tensorflow 1.5.0^[52]^, Keras 2.2.4^[53]^ and scikit-learn 0.19.1^[51]^ libraries. For each network, several variants in terms of number of layers and neurons, activation functions, regularization, loss functions and optimizer, were tested. In the end, we decided for two specific architectures that best fit our purpose: one deep FFNN trained on SS and one CNN trained on CBLV tree representation.

#### Deep feedforward neural network architecture for SS

The network consists of one input layer (of 99 input nodes both for trees with 50-199 and 200-500 tips), 4 sequential hidden layers organized in a funnel shape with 64-32-16-8 neurons and 1 output layer of size 2-4 depending on the number of parameters to be estimated. The neurons of the last hidden layer have linear activation, while others have exponential linear activation^[54]^.

| Layer (type) | Output Shape | Param # |
| --- | --- | --- |
| dense_1 (Dense) | (None, 64) | 6400 |
| dense_2 (Dense) | (None, 32) | 2080 |
| dense_3 (Dense) | (None, 16) | 528 |
| dense_4 (Dense) | (None, 8) | 136 |
| dense_5 (Dense) | (None, 2) | 18 |
| Total params: 9,162 |  |  |
| Trainable params: 9,162 |  |  |
| Non-trainable params: 0 |  |  |

##### Architecture: Feedforward neural network architecture

Example of FFNN trained on large trees to estimate the parameters of the BD model (R_0_ and infectious period 1/γ). ‘Dense’ layer means that for each neuron, all the inputs are multiplied by learned weights, summed together with the bias term. The activation function is then applied to the weighted sum before being output to the next layer. Dense_1 to dense_4 are layers with neurons of exponential linear activation, while dense_5 is composed either of softmax (in case of model selection) or of linear neurons (in case of parameter estimation). The number of trainable parameters in each layer is displayed (Param #): for example in the first layer, we have 99 input values and 1 bias for each of the 64 neurons, giving us in total (99+1)*64=6,400 trainable parameters. Output by Keras^[53]^, the ‘None’ in the ‘Output Shape’ means the network can input more than one training example at the time and that there is no constraint on the batch size (hence ‘None’).

#### Deep convolutional neural network for CBLV

The CNN consists of one input layer (of 400 and 1002 input nodes for trees with 50-199 and 200-500 tips, respectively). This input is then reshaped into a matrix of size of 201*2 and 501*2, for small and large trees, respectively, with entries corresponding to tips and internal nodes separated into two different rows (and one extra column with one entry in each row corresponding to the sampling probability). Then, there are two 1D convolutional layers of 50 kernels each, of size 3 and 10, respectively, followed by max pooling of size 10 and another 1D convolutional layer of 80 kernels of size 10. After the last convolutional layer, there is a GlobalPoolingAverage1D layer and a FFNN of funnel shape (64-32-16-8 neurons) with the same architecture and setting as the NN used with SS.

#### Neural network setting and training

For both NNs, we use the Adam optimisation algorithm^[55]^ as optimizer and the Mean Absolute Percentage Error (MAPE) as loss function. The batch size is set to 8,000. To train the network, we split the simulated dataset into 2 groups: ^[1]^ proper training set (3,990,000 examples); ^[2]^ validation set (10,000).

#### Preventing overfitting: Early stopping and Dropout

To prevent overfitting during training, we use: ^[1]^ the early stopping algorithm evaluating MAPE on a validation set; and ^[2]^ dropout that we set to 0.5 in the feed-forward part of both NNs^[56]^ (0.4, 0.45, 0.55 and 0.6 values were tried for basic BD model without improving the accuracy).

#### Neural networks for model selection

For model selection, we use the same architecture for FFNN-SS and CNN-CBLV as those for parameter inference described above. The only differences are: ^[1]^ the cost function: categorical cross entropy and ^[2]^ the activation function used for the output layer, that is, softmax function (of size 2 for small trees, selecting between BD and BDEI model, and of size 3 for large trees, selecting between BD, BDEI and BDSS). As we use the softmax function, the outputs of prediction are the estimated probabilities of each model, summing to 1.

The FFNN-SS and CNN-CBLV are trained on 8*10^6^ trees in the small tree setting (4*10^6^ trees per model, BD and BDEI). In the large tree setting, the FFNN-SS is trained on 12*10^6^ trees (4*10^6^ trees per model, BD, BDEI and BDSS and the CNN-CBLV is trained on 9*10^6^ trees (3*10^6^ trees per model, BD, BDEI and BDSS), instead of 12*10^6^ for GPU limitation purposes.

#### Predicting from very large trees using subtree picking and averaging

To predict from very large trees (*e.g.*, our ‘huge’ trees having 5,000 to 10,000 tips, **Fig. 4**) we designed the ‘Subtree Picker’ algorithm. The goal of Subtree Picker is to extract subtrees of bounded size representing independent sub-epidemics within the epidemic represented by the initial huge tree T, while covering most of the initial tree branches and tips in T. The sub-epidemics should follow the same sampling scheme as the global epidemic. This means that we can stop the sampling earlier than the most recent tip in T, but we cannot omit tips sampled before the end the sampling period (this would correspond to lower sampling probability). Each picked subtree corresponds to a sub-epidemic that starts with its root individual and lasts between its root date D_root_ and some later date (D_last_ > D_root_). The picked subtree corresponds to the top part of the initial tree’s clade with the same root, while the tips sampled after D_last_ are pruned.

The picked subtrees do not intersect with each other and contain between *m* and *M* tips each. Together they cover most of the initial tree’s branches. The initial tree T contains more than *M* tips. In the current PhyloDeep setting, *M*=500 (the largest tree size in the training set) and *m*=200 for BDSS and =50 for BD and BDEI (the smallest tree size in the training set).

Subtree Picker performs a postorder tree traversal (tips-to-root), where for each tree node N it calculates the maximum number of tips tN that can be extracted from its subtrees. The algorithm is recursive and combines two basic strategies: (1) the subtree rooted with N is decomposed into two independent sub-epidemics corresponding to N’s direct descendants; or (2) we decide to keep the upper part (with *x* oldest tips, *m* ≤ *x* ≤ M) of the subtree rooted with N, and the rest of N’s descendants is decomposed into independent sub-epidemics. The recursion is as follows (size(N) is the number of tips in the subtree rooted with N):

If size(N) < *m*, then t_N_=0

Else if *m* ≤ size(N) < *M* + *m*, then t_N_=max(size(N), M) #*pick the root subtree containing* t_N_ *oldest tips*

Else pick the best between the two strategies:

(1) Left L and right R children of N lead to independent sub-epidemics (subtrees): t_N_=t_L_+t_R_
(2) Pick the root subtree of *x* oldest tips (*m* ≤ *x* ≤ M, all possible *x* compared) plus the set Δ of the oldest descendant nodes of N, which represents the roots of independent sub-epidemics sampled after *x*^th^ tip date: t_N_=*x*+∑_D in Δ_ t_D_

For each processed node N, its optimal subtree picking strategy is memorized. Once t_N_ is calculated for the root of the global tree T, the algorithm picks the root’s subtrees according to the chosen strategy and, if needed, descends to the non-affected descendant nodes to pick more subtrees.

Let *s* be the size of T (number of tips). This tree decomposition requires one preorder tree traversal, with computing time in O(*s*). However, the picking strategy (2) requires another O(*s*) time to extract Δ (with appropriate data structure and pre-treatments). Thus, the whole computing time (computing (2) for each node in the tree traversal) is in O(*s*^2^), in the worst case. In practice, the subtrees extracted by Subtree Picker cover on average 98.5% (BD), 97.3% (BDEI) and 82.4% (BDSS) of the initial tree branches on the ‘huge’ tree datasets (5,000 to 10,000 tips). For the BDSS model this percentage is lower than for BD and BDEI, because of the narrower subtree size interval (*m*=200, *M*=500 versus *m*=50, *M*=500) corresponding to current PhyloDeep training set settings. In terms of computing time, Subtree Picker takes on average 0.6 (BD), 0.8 (BDEI) and 0.8 (BDSS) seconds per ‘huge’ tree, meaning that it could easily be applied to much larger trees.

Once subtrees (sub-epidemics) have been extracted, they are analysed using CNN-CBLV or FFNN-SS, and the parameter estimates are averaged with weights proportional to subtree sizes (number of tips).

### CONFIDENCE INTERVALS (95% CI)

#### Computation of 95% CI

We compute 95% CI using parametric bootstrap. To facilitate the deployment and speed-up the computation, we perform an approximation using a separate set of 1,000,000 simulations for calculation of CI. For each simulation in the CI set, we store the true parameter values *(i.e*., values with which we simulated the tree) and the parameter values predicted with both of our methods. This large dataset of true/predicted values is used to avoid new simulations, as required with the standard parametric bootstrap.

For a given simulated or empirical tree T, we obtain a set of predicted parameter values, {*p*}. The CI computation procedure searches among stored data those that are closest to T in terms of tree size, sampling probability and predicted values. We first subset:

- 10% of simulations within the CI set, which are closest to T in terms of size (number of tips), thus obtaining 100,000 CI sets of true/predicted parameter values.
- Amongst these, 10% of simulations that are closest to T in terms of sampling probability.

We thus obtain 10,000 CI sets of real/predicted parameter values, similar in size and sampling probability to T. For each parameter value *p* predicted from T, we identify the 1,000 nearest neighbouring values amongst the 10,000 true values of the same parameter available in the CI sets, 𝑅𝐶I={𝑟_𝑖=1,1000_}, and keep the corresponding predicted values, *P_CI_* = {𝑝𝑖=1,1000}. We then measure the errors for these nearest neighbors as 𝐸*_CI_*= {𝑒_𝑖_ = 𝑝_𝑖_ − 𝑟_𝑖_}. We center these errors around *p*, using the median of errors, *𝑚edian* (𝐸*_CI_*), which yields the distribution of errors for given prediction 𝑝: 𝐷 = {𝑝 + 𝑒_𝑖_ − *𝑚edian* (*E_CI_*)}, from which we extract the 95% CI around *p*. Individual points in the obtained distribution that are outside of the parameter ranges covered through simulations are set to the closest boundary value of the parameter range. For example, for f_SS_, if for a point in the distribution we obtain a value lower than 0.05, we set the value of that point to 0.05; and if we obtain a value larger than 0.20, we set it to 0.20. We undertake this procedure for all parameters except for the time related ones, that is, infectious and incubation period as these depend on the time rescaling. The width of our 95% CIs is defined as the distance between the 2.5% and 97.5% percentile. With very large trees and the subtree picking and averaging procedure, we consistently use a quadratic weighted average of the individual CIs found for every subtree.

#### Assessment of 95% CI coverage and width

To assess this fast implementation of the parametric bootstrap, we used the test set of 10,000 simulations (and 100 simulations for comparison with BEAST2 95% CI). We measured the coverage being defined as the fraction of simulations where the true/target parameter values are inside the obtained 95% CI:

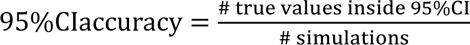

We applied the same criteria for BEAST2. For comparison of all methods, we excluded BDEI and BDSS simulations for which BEAST2 did not converge after 10 million steps. To draw BEAST2 CIs, we discarded the burn-in, that is, the first 10% of the MCMC, and calculated the CI on the remaining part of the chain. The CI width and coverage within the CIs obtained by NNs and BEAST2 are reported in **Supplementary Tab. 7**.

There exists a plethora of approaches for assessment of uncertainty and CI estimation. For example, (1) in a similar ABC context, the use of neighbouring trees (based on the Euclidean distance, not applicable to CBLV and questionable with SS) combined with a regression-based correction similar to that explained above^[19, 20]^; (2) the (non-approximated) parametric bootstrap^[56]^; ^[3]^ the prediction of values from a distribution of trees reconstructed with Bayesian methods^[10]^; *etc*. We chose an approximation of the parametric bootstrap for its easy deployability, speed, coverage and width of produced CIs. The easy deployability comes from the fact that CIs are based on pre-calculated data stored in our CI set. The speed of the method comes from it not requiring simulations of new trees, and thus producing CIs within 2-4 seconds. The coverage and width are comparable to those of BEAST2 (**Sup. Table 4**), a Bayesian method, intended to estimate the distribution of parameters and the uncertainty of inferences, with high computational cost.

### MODEL ADEQUACY

#### A priori checks

We performed a sanity check using the SS of the test set simulations and the SS measured on the empirical HIV phylogeny. We reduced and centered the SS and performed a Principal Component Analysis (PCA) using the PCA function from the scikit-learn^[51]^ package.

We highlighted the data point corresponding to the Zürich HIV MSM phylogeny in **Supplementary Fig. 8**, for each model (BD, BDEI and BDSS). Dissemblance between the simulations and the HIV phylogeny would be manifested by the fact that this data point lies outside the distribution corresponding to the simulations.

Furthermore, we performed an additional *a priori* check consisting in the study of all individual SS rather than dimensional reduction with PCA. For each SS, we checked whether the value for Zürich HIV MSM phylogeny lays between the minimum and maximum value of that SS in the test set of 10.000 trees. We reported the results in **Supplementary Fig. 8**.

#### A posteriori checks

We performed tests analogous to the *a priori* model adequacy checks. For both PCA and individual SS tests, instead of using the test set as representative of simulations, we simulated 10,000 additional simulations under the selected BDSS model. Parameter values were resampled from uniform distribution with boundaries given by the 95% CIs, and sampling probability fixed to presumed value of 0.25 (**Fig. 5, Supplementary Fig. 8**).

### MODELS

The models we used for tree simulations are represented in the form of flow diagrams in **Fig. 1**. We simulated dated binary trees for (1) the training of NNs and (2) accuracy assessment of parameter estimation and model selection. We used the following three individual-based phylodynamic models:

#### Constant rate birth-death model with incomplete sampling

This model (BD^[8, 9]^, **Fig. 1 a**) contains three parameters and three compartments: infectious (I), removed with sampling (R) and removed unsampled (U) individuals. Infection takes place at rate β. Infectious individuals are removed with rate γ. Upon removal, an individual is sampled with probability *s*.

For simulations, we re-parameterized the model in terms of: basic reproduction number, R_0_; infectious period, 1/γ; sampling probability, *s*; and tree size, *t.* We then sampled the values for each simulation uniformly at random in the ranges given in **Supplementary Tab. 4**.

#### Birth-death model with exposed-infectious classes

This model (BDEI^[10–12]^, **Fig. 1 b**) is a BD model extended through the presence of an exposed class. More specifically, this means that each infected individual starts as non-infectious (E) and becomes infectious (I) at incubation rate ε. BDEI model thus has four parameters (β, γ, ε and *s*) and four compartments (E, I, R and U).

For simulations, we re-parameterized the model similarly as described for BD, set the ε value via 1/γ and incubation ratio (=ε/γ). We sampled all parameters, including ε/γ, from a uniform distribution, just as with BD (**Supplementary Tab. 4**).

#### Birth-death model with superspreading

This model (BDSS^[5,10,11]^, **Fig. 1 c**) accounts for heterogeneous infectious classes. Infected individuals belong to one of two infectious classes (I_S_ for superspreading and I_N_ for normal spreading) and can transmit the disease by giving birth to individuals of either class, with rates β_S,S_ and β_S,N_ for I_S_ transmitting to I_S_ and to I_N_, respectively, and β_N,S_ and β_N,N_ for I_N_ transmitting to I_S_ and I_N_, respectively. However, there is a restriction on parameter values: β_S,S_ ∗ β_N,N_ = β_S,N_ ∗ β_N,S_. There are thus superspreading transmission rates β_S,._ and normal transmission rates β_N,._ that are X_SS_ (= β_S,S_/β_N,S_ = β_S,N_/β_N,N_) times higher for superspreading. At transmission, the probability of the recipient to be superspreading is f_ss_ (= β_S,S_/(β_S,S_ + β_S,N_)), the fraction of superspreading individuals at equilibrium. We consider that both I_S_ and I_N_ populations are otherwise indistinguishable, that is, both populations share the same infectious period (1/γ)^[5,10,11]^. The model thus has six parameters, but only five need to be estimated to fully define the model^[5, 10]^.

For simulations, we chose parameters of epidemiological interest for re-parameterization: basic reproduction number (𝑅_0_), infectious period 1/γ, f_SS_, X_ss_ and sampling probability *s*. In our simulations, we used uniform distributions for these 5 parameters, just as with BD and BDEI (**Supplementary Tab. 4**).

### SIMULATIONS

For the parameters R_0,_ 1/γ, and *s*, that are common to all three birth-death models, the same value boundaries were used across all models (**Supplementary Tab. 4**). We considered two spans of tree size: ‘small trees’ with 50 to 199 tips and ‘large trees’ with 200 to 500 tips. We then sampled parameter values uniformly at random within these parameter boundaries with standard Latin-hypercube sampling^[57]^ using PyDOE package. We created 3,990,000 parameter sets for training, 10,000 for validation and early stopping, another 10,000 for testing parameter inference and model selection (comparison with BEAST2 used a subset of 100, for computing time reasons), and 1,000,000 parameter sets for fast computation of CIs.

With these parameter sets, we simulated trees under each birth-death model using our implementation in Python of Gillespie algorithm^[58]^, based on a standard forward simulator. Comparable accuracies (as in **Fig. 3 and Supplementary Fig. 2**, both for BEAST2 and our methods) were reached on test simulations obtained with a well-established (but slower) simulator: TreeSim^[4,5,7]^ (data not shown).

Each simulation started with one infectious individual (the class was chosen randomly under the BDSS model) and stopped when we obtained a tree with the given number of sampled individuals (tips). If the epidemic died away stochastically, that is, there was no more infectious tips left due to stochastic death before reaching the given tree size, we re-initialized the simulation up to 100 times. Only around 11% of simulations reached more than 2 iterations (20% for BDSS), and less than 0.5% reached more than 50 iterations for all models. If still no tree of given size was obtained after 100 iterations, we discarded the parameter set (less than 0.3% of all sets) and generated a new one to keep the desired number of simulations. This enabled us to maintain a nearly uniform coverage of parameter space, within selected parameter boundaries.

We simulated 100 ‘huge’ trees for each model, with the same parameter values as for the 100 large trees used for testing (Fig. 3). These trees were simulated with treesimulator (https://pypi.org/project/treesimulator). The Snakemake^[61]^ pipeline for huge trees simulation along with the simulated trees are available on GitHub.

### METHOD COMPARISON

#### Parameter inference with BEAST2

To assess the accuracy of our methods, we compared it with a well-established Bayesian method, as implemented in BEAST2 (version 2.6.2). We used the BDSKY package^[4]^ (version 1.4.5) to estimate the parameter values of BD simulations and the package bdmm^[12, 13]^ (version 1.0) to infer the parameter values of BDEI and BDSS. Furthermore, for the inference on BDSS simulations, instead of BEAST 2.6.2 we used the BEAST2 code up to the commit nr2311ba7, which includes important fixes to operators critical for our analyses. We set the Markov Chain Monte Carlo (MCMC) length to 5 million steps for the BD model, and to 10 million steps for the BDEI and BDSS models.

The sampling probability was fixed during the estimation. Since the BD, BDEI and BDSS models implemented in BEAST2 do not use the same parametrizations as our methods, we needed to apply parameter conversions for setting the priors for BEAST2 inference (**Supplementary Tab. 6**), and for translating the BEAST2 results back to parameterizations used in our methods, in order to enable proper comparison of the results. More specifically, the BEAST2 parameters can be converted to those used in our methods, that is, instead of infectious period and incubation period, BEAST2 uses the inverse of these, namely the infectious rate and incubation rate, respectively; instead of superspreading transmission ratio and superspreading fraction at equilibrium, it uses individual sub-component parameters R_0,SS_, R_0,SN_, R_0,NS_ and R_0,NN_, which we will collectively refer to as “partial R_0_”. For BDSS, the BEAST2 prior was thus not the same as that of our simulations for BDSS (**Supplementary Tab. 4, 6**), since BEAST2 does not infer the same parameters. We used the range of all parameter values used in our simulations to set the boundaries of uniform prior distributions of parameters inferred by BEAST2. The initial values in the MCMC were set to the medians observed in the training set. During the inference, the parameter values were constrained in the same way as in the simulations, namely, we used the following constraint 𝑅_0,𝑁𝑁_∗𝑅_0,𝑆𝑆_=𝑅_0,𝑆𝑁_∗𝑅_0,𝑁𝑆_(equivalent to 𝛽_𝑁,𝑁_∗𝛽_𝑆,𝑆_=𝛽_𝑆,𝑁_∗𝛽_𝑁,𝑆_) in the BDSS model inference. Furthermore, the effective frequency of superspreading individuals (parameter called “geo-frequencies” in *bdmm*) was constrained to be between 5% and 20%. Due to the parameter conversions, and despite these constraints the inferred f_ss_ and X_ss_ can reach values outside the boundaries used for simulations, in which case we set them to the closest boundary for fair comparison with deep learning methods in **Fig. 3** (*e.g.*, if the median *a posteriori* f_ss_ was estimated to be larger than 0.20, it was set to 0.20 and if inferred f_ss_ was less than 0.05, it was set to 0.05). The goal of this correction was to avoid penalizing BEAST2 when it converged to local minima outside of the parameter boundaries used for simulations, which are implicitly known to NNs since they were trained on simulations with parameters within these boundaries.

After we obtained the parameters of interest from the original parameters estimated by BEAST2, we evaluated the Effective Sample Size (ESS) on all parameters. We reported the absolute percentage error of the median of *a posteriori* values, corresponding to all reported steps (reported steps being spaced by 1,000 actual MCMC steps) past the 10% burn-in. For simulations for which BEAST2 did not converge, we considered the median of the parameter distribution used for simulations (**Fig. 3, Supplementary Tab. 1-2, Supplementary Fig. 2**) or excluded them from the comparison (**Supplementary Tab. 1-2**, values reported in brackets, **Supplementary Tab. 5**).

For the HIV application, the prior of infectious period was set to [0.1, 30] years (uniform). All the other parameters had the same prior distributions as used in simulations and shown in **Supplementary Tab. 4, 6.**

#### Model selection with BEAST2

We performed model selection under BEAST2 using Akaike’s information criterion through MCMC (AICM)^[32, 33]^. The AICM is based on the following formula:

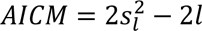

where *l* and 𝑠_j_^2^ are the sample mean and variance of the posterior log-likelihoods. The AICM is an equivalent of AIC and the model with lowest AICM value is selected.

For 100 simulations obtained with each model (BD, BDEI and BDSS for large trees, BD and BDEI for small trees), we performed parameter estimation with BEAST2 under each model, computed AICM considering the whole MCMC, but excluding 10% burn-in (*i.e.*, 9,000 log-likelihood values for BDEI and BDSS considered in total, 4,500 for BD). The results of model selection are shown in **Supplementary Tab. 5**. The BDEI and BDSS simulations for which BEAST2 did not reach an ESS of 200 for all parameters were excluded from the computation of model selection accuracy for all methods.

#### Linear regression

For each model, linear regression was trained using reduced and centered summary statistics (using scikit-learn package, as with FFNN). Its bias and accuracy were assessed using the same criteria as for the NN approaches (**Supplementary Tables 1-2, Supplementary Fig. 6**).

#### FFNN-CBLV

We trained an FFNN on CBLV representation. The FFNN architecture was close to the one described in ***Architecture*** with one extra hidden layer, so 5 layers in total, organized in a funnel shape with 128-64-32-16-8 neurons and 1 output layer of size 2-4 depending on the number of parameters to be estimated. The setting during the training and the sizes of training, validation and testing sets were the same as for the CNN-CBLV. Its bias and accuracy were assessed using the same criteria as for other NN approaches (**Supplementary Tab. 1-2, Supplementary Fig. 5**).

#### TreePar

We used TreePar^[5]^ for MLE. With BD, we obtained results close to estimates under BEAST2, which is consistent with former studies^[58]^. TreePar^[5]^ uses an exact analytical formula of likelihood for BD and thus these (and BEAST2) results are theoretically optimal.

We also performed several trials to do parameter inference for the more complex models (*i.e.*, BDEI and BDSS), but in a large number of cases, we encountered numerical problems (*e.g.*, underflow or overflow issues), which resulted in infinite negative log-likelihood values, and eventually failed runs. When the calculations did not fail, we found that

many estimations under BDSS and BDEI had lower likelihood than estimations performed with (nested) BD on the same input data. These numerical issues were confirmed by the authors of the TreePar package, with no solution available at the moment.

#### Null models

To assess how much information was learned on given problem, we compared FFNN-SS and CNN-CBLV to two null models.

The first null model was the FFNN trained for each model on 4,000,000 simulations using SS, but with randomly permuted target values (*i.e*., the initial correspondence between the SS and underlying parameter values was lost, while the range of values was conserved). We then predicted parameters for 10,000 test simulations (100 for comparison with BEAST2) and measured the mean absolute relative error (MRE; **Supplementary Tab. 1**). In such a case, the FFNN always predicted values close to the value with the lowest value of the cost function (*e.g.*, 2.2 for parameter values uniformly sampled between 1 and 5). The MRE of this approach represents the lowest MRE that machine learning approaches can have in the absence of information, but the knowledge of the parameter distribution. This can be used to get an idea of how well the trained approaches perform and how much information regarding each parameter they can extract from the data.

The second null model was a set of random values sampled from the parameter ranges that were used for simulations (**Supplementary Tab. 4**). In this model, as opposed to the previous null model, there is no training phase and we do not learn the best compromise in the absence of information.

### PERFORMANCE ASSESSMENT

#### Mean relative error MRE

To compare the accuracy of parameter estimation, we used 100 simulated trees per model. We computed the mean absolute relative error (MRE, **Fig. 3-4, Supplementary Tab. 1, Supplementary Fig. 2, 4**) between (1) the true (or target) parameter values and the predicted values for machine learning approaches; and (2) the true (or target) parameter values and the median *a posteriori* values obtained with BEAST2, which are more stable and accurate than maximum *a posteriori* values:

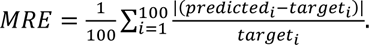

We plotted individual absolute relative errors (RE) of predictions (**Fig. 3-4, Supplementary Fig. 2, 4**) for each simulation *i*, calculated as:

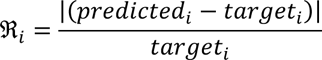

Not being limited by the computational cost for machine learning approaches, we computed the same metric but on 10,000 simulations (**Supplementary Fig. 3, 5-6;** results from 1,000 simulations plotted in **Supplementary Fig. 7**).

We assessed the statistical significance of MRE differences using paired z-test. The two NN approaches were also compared using the same test, but no significant differences were found.

#### Mean relative bias MRB

To compare the bias in parameter estimation, we used 100 simulated trees per model. We computed the mean relative bias (MRB) between ^[1]^ the true (or target) parameter values and the predicted values for machine learning approaches; and ^[2]^ the true (or target) parameter values and the median *a posteriori* values obtained with BEAST2 (**Supplementary Tab. 2**):

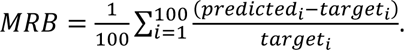

#### Comparison of likelihood values for sets of parameter estimates obtained with different methods

To assess the performance of different methods, we also studied the likelihood values of parameter estimates obtained with BEAST2, CNN-CBLV and FFNN-SS. For BD, we computed the likelihood using TreePar and compared it to the likelihood value of target parameter values (**Supplementary Tab. 3**).

As TreePar was problematic with BDEI and BDSS (see above), we tried to take on the same approach for BDEI and BDSS with BEAST2, but imposing a single MCMC step. Nevertheless, this did not yield sufficient results to perform sound comparison, since for example with FFNN-SS predictions, the likelihood was obtained only for 57/100 parameter estimates for BDEI and 49/100 for BDSS. In the remaining cases, BEAST2 either failed to return consistent likelihood values, or was unable to calculate likelihood for the initial parameter values.

#### Model selection accuracy

We performed model selection with CNN-CBLV, FFNN-SS and BEAST2 on 100 simulations obtained with each model (10,000 for a sub-comparison of CNN-CBLV and FFNN-SS). Results are shown in **Supplementary Tab. 5** in the form of confusion matrices, where the columns represent the true/target classes, and the rows are the predicted classes.

We then computed the accuracy of each method:

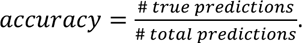

For BEAST2 model selection and large trees, the chain did not converge (displayed as “ESS<200” in **Supplementary Tab. 5**) for 24.3% simulations of large trees and 4.5% simulations of small trees. We did not consider these in accuracy measurements, for all the methods.

#### Comparison of time efficiency

For FFNN-SS and CNN-CBLV, we reported the average CPU time of encoding a tree (average over 10,000 trees), as reported by NextFlow workflow manager^[60]^, a pipeline software that we used. The inference time itself was negligible.

For BEAST2, we reported the CPU time averaged over 100 analyses with BEAST2 as reported by NextFlow. For the analyses with BDEI and BDSS models, we reported the CPU time to process 10 million MCMC steps, and for the analyses with BD, we reported the CPU time to process 5 million MCMC steps. To account for convergence, we re-calculated the average CPU time considering only those analyses for which the chain converged and an ESS of 200 was reached across all inferred parameters.

The calculations were performed on a computational cluster with CentOS machines and Slurm workload manager. The machines had the following characteristics: 28 cores, 2.4Ghz, 128 GB of RAM. Each of our jobs (simulation of one tree, tree encoding, BEAST2 run, etc.) was performed requesting one CPU core. The neural network training was performed on a GPU cluster with Nvidia Titan X GPUs.

### HIV DATASET

We used the original phylogenetic tree reconstructed by Rasmussen *et al.*^[25]^ from 200 sequences corresponding to the largest cluster of HIV-infected men-having-sex-with-men (MSM) subpopulation in Zurich, collected as a part of the Swiss Cohort Study^[24]^. For details on tree reconstruction, please refer to their article.

### PHYLODEEP SOFTWARE

FFNN-SS and CNN-CBLV parameter inference, model selection, 95% CI computation and *a priori* checks are implemented in the PhyloDeep software, which is available on GitHub (github.com/evolbioinfo/phylodeep), PyPi (pypi.org/project/phylodeep) and Docker Hub (hub.docker.com/r/evolbioinfo/phylodeep). It can be run as a command-line program, Python3 package and a Docker container. PhyloDeep covers the parameter subspace as described in **Supplementary Tab. 4**. The input is a dated phylogenetic tree with at least 50 tips and presumed sampling probability. The output is a PCA plot for *a priori* check, a csv file with all SS and their bounds in simulated data for the three models, and a csv file with probabilities of each model (for model selection) and point estimates and 95% CI values (for parameter inference with selected model). The installation details and usage examples are available as well on GitHub.

### DATA AND CODE AVAILABILITY

Together with the PhyloDeep package source code, we provide on GitHub (github.com/evolbioinfo/phylodeep): (i) all data shown in the article, (ii) the code of the tree simulators used to train the deep learners, (iii) the HIV phylogeny analysed in the article as a showcase application.

