## Supplementary material for "Deep learning from phylogenies to uncover the epidemiological dynamics of outbreaks": Voznica_et_al_Supplementary_information

**Page**

#### TABLES

**Supplementary Table 1: Method comparison in terms of accuracy** 2

**Supplementary Table 2: Method comparison in terms of bias** 3

**Supplementary Table 3: Method comparison in terms of likelihood with BD model** 4

**Supplementary Table 4: Parameter ranges used for simulations** 5

**Supplementary Table 5: Method comparison in terms of model selection** 6

**Supplementary Table 6: BEAST2 parameters and priors** 7

**Supplementary Table 7: Confidence interval assessment** 8

#### FIGURES

**Supplementary Figure 1: Bijectivity of the CBLV representation** 9

**Supplementary Figure 2: Assessment of deep learning accuracy on small trees** 10

**Supplementary Figure 3 Accuracy of deep learning methods increases with tree size** 11

**Supplementary Figure 4: Assessment of deep learning generalization capabilities** 12

**Supplementary Figure 5: CNN performs well with CBLV tree representation** 13

**Supplementary Figure 6: How much and how fast FFNN-SS and CNN-CBLV learn?** 14

**Supplementary Figure 7: Adding new SS to increase accuracy of FFNN-SS** 16

**Supplementary Figure 8: *A priori* and *a posteriori* checks of model adequacy for HIV data** 17

**Supplementary Table 1: Method comparison in terms of accuracy**

| Model | Parameter | Mean Relative Error | | | | | | | |
|---|---|---|---|---|---|---|---|---|---|
| | | 1. FFNN SS | 2. CNN CBLV | 3. BEAST2 | 4. FFNN CBLV | 5. LR SS | 6. Null model 1 | 7. Null model 2 | z-test (<0.05) |
| **BD 200-500 tips** | $R_0$ | **0.07** | **0.07** | **0.06** | **0.08** | 0.19 | 0.36 | 0.56 | 1=2=3=4 |
| | $1/\gamma$ | **0.06** | **0.06** | **0.06** | **0.06** | 0.22 | 0.30 | 0.82 | 1=2=3=4 |
| **BDEI 200-500 tips** | $R_0$ | **0.08** (0.08) | **0.09** (0.09) | **0.10** (0.10) | **0.10** (0.10) | 0.20 (0.20) | 0.43 (0.43) | 0.56 | 1=2=3=4 |
| | $1/\varepsilon$ | **0.13** (0.13) | **0.13** (0.13) | 0.20 (0.19) | 0.29 (0.29) | 0.21 (0.21) | 0.74 (0.74) | 2.08 | 1=2 |
| | $1/\gamma$ | **0.10** (0.10) | **0.10** (0.10) | 0.31 (0.30) | 0.20 (0.21) | 0.26 (0.26) | 0.51 (0.51) | 0.82 | 1=2 |
| **BDSS 200-500 tips** | $R_0$ | **0.09** (0.09) | **0.10** (0.10) | 0.16 (0.11) | **0.10** (0.10) | 0.18 (0.17) | 0.34 (0.35) | 0.56 | 1=2=4 |
| | $1/\gamma$ | **0.09** (0.09) | **0.09** (0.09) | 0.22 (0.10) | **0.10** (0.10) | 0.18 (0.16) | 0.26 (0.27) | 0.84 | 1=2=4 |
| | $X_{SS}$ | **0.11** (0.11) | **0.13** (0.13) | 0.19 (0.14) | 0.23 (0.22) | **0.15** (0.14) | 0.27 (0.28) | 0.41 | 1=2=5 |
| | $f_{SS}$ | **0.25** (0.23) | **0.26** (0.23) | 0.43 (0.44) | 0.33 (0.33) | 0.36 (0.36) | 0.34 (0.34) | 0.47 | 1=2 |

For each model and inferred parameter, we compared the different inference methods in terms of Mean Relative Error (). We compared these measures for 1. FFNN trained on SS, 2. CNN trained on CBLV, 3. BEAST2, 4. FFNN trained on CBLV, 5. Linear regression trained on SS, 6. "Null model 1", *i.e.* FFNN trained on SS with permuted target values and 7. "Null model 2" measured on 100 targets, where for each target we sampled randomly values from the prior parameter subspace. For 1.-6. MRE values were measured on the same 100 simulations. For BEAST2, we considered median values of covered parameter subspace, when BEAST2 did not converge (2% and 15% of simulations for BDEI and BDSS, respectively). For 1.-6. we displayed in parentheses the MRE when not considering the simulations for which BEAST2 did not converge. We compared the individual methods (1.-6.) in a pairwise fashion to find out if one was more accurate in terms of MRE than the other, using paired z-test, at significance level of 0.05 and we show the most accurate methods for each parameter in bold.

In all settings, FFNN-SS and CNN-CBLV are amongst the most accurate methods, being exclusively the most accurate ones for BDEI $1/\gamma$ and $1/\varepsilon$ and for BDSS $f_{SS}$. Furthermore, the comparison of MRE values shows that the prediction of $f_{SS}$ has low accuracy for all methods (0.25 and 0.26 for FFNN-SS and CNN-CBLV, respectively, vs 0.34 obtained with "Null model 1"). This might be due to low information on superspreading individuals in the trees of this size, as supported by Supplementary Fig. 3.

**Supplementary Table 2: Method comparison in terms of bias**

| Model | Parameter | Mean Relative Bias | | | | |
|---|---|---|---|---|---|---|
| | | 1. FFNN SS | 2. CNN CBLV | 3. BEAST2 | 4. FFNN CBLV | 5. LR SS |
| **BD 200-500 tips** | $R_0$ | -0.01 | -0.01 | -0.01 | -0.01 | 0.04 |
| | $1/\gamma$ | -0.01 | -0.02 | -0.02 | -0.01 | 0.04 |
| **BDEI 200-500 tips** | $R_0$ | -0.03 (-0.03) | -0.03 (-0.03) | -0.04 (-0.04) | 0.04 (0.04) | -0.03 (-0.03) |
| | $1/\varepsilon$ | -0.01 (-0.01) | -0.01 (-0.01) | **-0.10 (-0.12)** | 0.02 (0.03) | 0.02 (0.02) |
| | $1/\gamma$ | -0.03 (-0.03) | -0.01 (-0.01) | **0.24 (0.23)** | -0.03 (-0.03) | -0.01 (-0.01) |
| **BDSS 200-500 tips** | $R_0$ | -0.01 (-0.01) | -0.01 (-0.01) | **0.07** (0.04) | 0.00 (-0.01) | 0.02 (0.00) |
| | $1/\gamma$ | 0.00 (0.00) | 0.00 (-0.01) | **0.12** (0.03) | 0.00 (-0.01) | 0.03 (0.00) |
| | $X_{SS}$ | -0.01 (-0.02) | 0.01 (0.00) | **0.05** (-0.02) | **0.07** (0.04) | **0.05** (0.03) |
| | $f_{SS}$ | -0.03 (-0.01) | 0.02 (0.03) | **0.32 (0.36)** | -0.02 (-0.01) | **0.19 (0.20)** |

For each model and inferred parameter, we compared the different inference methods in terms of Mean Relative Bias (MRB). We compared the MRB for 1. FFNN trained on SS, 2. CNN trained on CBLV, 3. BEAST2, 4. FFNN trained on CBLV and 5. Linear regression trained on SS. These MRB values were measured on the same 100 simulations. If BEAST2 did not converge, we set the inferred value to the median of covered parameter subspace. We further display MRB when not considering simulations for which BEAST2 did not converge, shown in parentheses. The biases bigger than 0.05 are displayed in red bold.

This shows that FFNN-SS and CNN-CBLV have low bias, while BEAST2 suffers from an intermediate to high bias with BDEI and BDSS for most parameters. These biases may account for a large fraction of MRE for BEAST2.

**Supplementary Table 3: Method comparison in terms of likelihood with BD model**

| Y > X | Y: True values | Y: BEAST2 | Y: CNN-CBLV | Y: FFNN-SS |
|---|---|---|---|---|
| X: True values | - | 70 | 66 | 69 |
| X: BEAST2 | 30 | - | 43 | 53 |
| X: CNN-CBLV | 34 | 57 | - | 55 |
| X: FFNN-CBLV | 31 | 47 | 45 | - |

| Median Y-X | Y: True values | Y: BEAST2 | Y: CNN-CBLV | Y: FFNN-SS |
|---|---|---|---|---|
| X: True values | 0.0 | 4.9 | 4.4 | 4.5 |
| X: BEAST2 | - | 0.0 | -0.2 | 0.2 |
| X: CNN-CBLV | - | - | 0.0 | 0.2 |
| X: FFNN-CBLV | - | - | - | 0.0 |

For BD model and 100 large test trees (200-500 tips), we compared parameter estimates obtained with different inference methods in terms of likelihood. We compared loglikelihood values evaluated by TreePar package for: True values with which test trees were simulated; estimates obtained with BEAST2; estimates obtained with CNN-CBLV; estimates obtained with FFNN-SS. The first table shows a simple pairwise comparison (e.g. 30 means that the True value was better than BEAST2 in 30% of cases), while the second table shows the median of differences between loglikelihood values.

The two results go in the same direction. The likelihood of both FFNN-SS and CNN-CBLV estimates is similar to BEAST2's, which explains the similar accuracy of the three methods (**Fig. 3**). Regarding the comparison with 'True values', if a given method tends to produce higher likelihood than that of the true parameter values, then it performs well in terms of likelihood optimization, as optimizing further should not result in higher accuracy. The results are again quite positive, as BEAST2 as well as our NNs achieved a higher likelihood than the true parameter values for ~70% of the trees, with a significant mean difference.

**Supplementary Table 4: Parameter ranges used for simulations**

| Parameters | Name | Range | Inferred/set | Relation to other parameters |
|---|---|---|---|---|
| $R_0$ | basic reproduction number | U(1,5) | inferred | $= \beta/\gamma$ (BD & BDEI)<br>$= (\beta_{S,S} + \beta_{N,N})/\gamma$ (BDSS) |
| $1/\gamma$ | infectious period | U(1,10) | inferred | |
| t | tree size | "small" trees: U(50, 199);<br>"large" trees: U(200, 500) | set | |
| s | sampling probability | U(0.01,1) | set | |
| $f_I$ | incubation factor | U(0.2,5) | parameterization | $= \varepsilon/\gamma$ |
| $1/\varepsilon$ | incubation period | [0.2, 50] | inferred | $= 1/(f_I{*}\gamma)$ |
| $X_{SS}$ | superspreading infectious ratio at equilibrium | U(3,10) | inferred | $= \beta_{S,S}/\beta_{N,S} = \beta_{S,N}/\beta_{N,N}$ |
| $f_{SS}$ | fraction of superspreading individuals at equilibrium | U(0.05, 0.20) | inferred | $= \beta_{S,S}/(\beta_{S,S}+\beta_{S,N})$<br>$= 1-\beta_{N,N}/(\beta_{N,N}+\beta_{N,S})$ |

For each parameter, we display its full name and the parameter range covered by simulations of training and testing sets. Note that all parameters were sampled from a uniform distribution, which we denote by U(x,y), with x being the lower and y the upper bound of that uniform distribution. The table further shows whether these parameters were inferred or used as input and, where appropriate, their relation to other model parameters (displayed in Figure 1). Parameters common to all three models are coloured in yellow, BDEI-specific in purple and BDSS-specific in green. The models are parameterized by the parameters of epidemiological interest except for the incubation factor, which enables to set the incubation period to values that are reasonable with respect to the infectious period. More specifically, through our parameterization choices, the incubation factor spans from 20% to 500% of the value of the infectious period, making it both not negligible but also not too high when compared to infectious period.

**Supplementary Table 5: Method comparison in terms of model selection**

**a (i)**

| FFNN-SS | Actually BD | Actually BDEI |
|---|---|---|
| Predicted BD | 94 (9360) | 13 (1837) |
| Predicted BDEI | 6 (640) | 87 (8163) |

**a (ii)**

| FFNN-SS | Actually BD | Actually BDEI | Actually BDSS |
|---|---|---|---|
| Predicted BD | 93 (9488) | 11 (1006) | 4 (561) |
| Predicted BDEI | 6 (304) | 86 (8783) | 0 (281) |
| Predicted BDSS | 1 (208) | 3 (211) | 96 (9158) |

**b (i)**

| CNN-CBLV | Actually BD | Actually BDEI |
|---|---|---|
| Predicted BD | 96 (9289) | 17 (1970) |
| Predicted BDEI | 4 (711) | 83 (8030) |

**b (ii)**

| CNN-CBLV | Actually BD | Actually BDEI | Actually BDSS |
|---|---|---|---|
| Predicted BD | 91 (9131) | 10 (1052) | 3 (514) |
| Predicted BDEI | 7 (519) | 88 (8781) | 1 (137) |
| Predicted BDSS | 2 (350) | 2 (167) | 96 (9349) |

**c (i)**

| BEAST2 | Actually BD | Actually BDEI |
|---|---|---|
| Predicted BD | 75 | 4 |
| Predicted BDEI | 21 | 91 |
| ESS<200 | 4 | 5 |

**c (ii)**

| BEAST2 | Actually BD | Actually BDEI | Actually BDSS |
|---|---|---|---|
| Predicted BD | 62 | 0 | 2 |
| Predicted BDEI | 6 | 72 | 0 |
| Predicted BDSS | 6 | 7 | 72 |
| ESS<200 | 26 | 21 | 26 |

Confusion matrices obtained with **a (i-ii)**, FFNN-SS, **b (i-ii)**, CNN-CBLV and **c (i-ii),** BEAST2 using AICM, either with **a-c (i)**, small trees (between 50 and 199 tips) or **a-c (ii)**, large trees (between 200 and 500 tips). BDSS and BDEI are nested within BD, namely, BDEI becomes BD when incubation period is 0 and BDSS becomes BD when superspreading individuals transmit at the same rate as normal individuals. We display the results as confusion matrices, actual classes being columns and predicted ones being rows. These were obtained with a test set of 100 simulations obtained with each model (or 10,000 for FFNN-SS and CNN-CBLV, results in parentheses). For BEAST2, 76% simulations converged for large trees and 96% for small trees. We show in red the number of simulations that did not reach an ESS of 200 for at least one parameter.

**Supplementary Table 6: BEAST2 parameters and priors**

| Parameters | Name | BEAST2 parameter | Range | Initial value | Formula |
|---|---|---|---|---|---|
| γ | become uninfectious rate | Yes | U(0.1,1.0) | 0.55 | |
| 1/γ | infectious period | No | [1,10] | 1.8 | |
| s | sampling probability | Yes | fixed | true value | |
| $R_0$ | basic reproduction number | Yes | U(1.0,5.0) | 3.0 | = β/γ (BD & BDEI) |
| ε | incubation rate | Yes | U(0.02,5.0) | 2.51 | |
| 1/ε | incubation period | No | [0.2,50] | 0.40 | |
| $R_{0,SS}$ | partial, within deme superspreading $R_0$ | Yes | U(0.14,4.31) | 1.25 | |
| $R_{0,NN}$ | partial, within deme normal spreading $R_0$ | Yes | U(0.14,4.31) | 1.44 | |
| $R_{0,SN}$ | partial, outside deme superspreading $R_0$ | Yes | U(0.034,32.30) | 9.0 | |
| $R_{0,NS}$ | partial, outside deme normal spreading $R_0$ | Yes | U(0.034,32.30) | 0.20 | |
| $R_0$ | basic reproduction number | No | [0.28,8.62] | 2.69 | $= R_{0,SS} + R_{0,NN}$ (BDSS) |
| $X_{SS}$ | superspreading infectious ratio at equilibrium | No | [4*10$^{-4}$,127] | 6.25 | $= R_{0,SS}/R_{0,NS} = R_{0,SN}/R_{0,NN}$ |
| $f_{SS}$ | fraction of superspreading individuals at equilibrium | No | [4*10$^{-3}$,0.99] | 0.12 | $= R_{0,SS}/(R_{0,SS}+R_{0,SN})$<br>$= 1-R_{0,NN}/(R_{0,NN}+R_{0,NS})$ |

This table shows parameters and their prior distributions used during inference with BEAST2. We display the parameters, their definitions and priors in BEAST2, that are common to all models (in yellow), common to BD and BDEI (in red), BDEI-specific (in purple) and BDSS-specific (in green). From these parameters, we deduce the values and distributions of parameters of interest as shown in the table. Note that the parameters of epidemiological interest are basic reproduction number and infectious period for BD, BDEI and BDSS, incubation period for BDEI, and superspreading infectious ratio and fraction of superspreading individuals for BDSS. We check convergence (ESS) on all parameters and extract median *a posteriori* and CI values exclusively for the parameters of epidemiological interest.

**Supplementary Table 7: Confidence interval assessment**

| Model | Parameter | Range | 95% CI width and coverage | | | | | |
|---|---|---|---|---|---|---|---|---|
| | | | FFNN-SS computed on converged simulations | | CNN-CBLV computed on converged simulations | | BEAST2 computed on converged simulations | |
| | | | Coverage | Width | Coverage | Width | Coverage | Width |
| BD 200-500 tips | $R_0$ | U(1, 5) | 0.99 (0.93) | 0.99 (1.0) | 0.98 (0.93) | 1.0 (1.1) | 0.99 | 0.99 |
| | $1/\gamma$ | U(1, 10) | 0.97 (0.92) | 1.6 (1.5) | 0.97 (0.92) | 1.6 (1.6) | 1.00 | 1.6 |
| BDEI 200-500 tips | $R_0$ | U(1, 5) | 0.89 (0.93) | 1.0 (1.1) | 0.92 (0.93) | 1.1 (1.1) | 0.88 | 1.0 |
| | $1/\varepsilon$ | [0.2, 50] | 0.89 (0.91) | 6.6 (6.6) | 0.88 (0.92) | 7.1 (7.2) | 0.84 | 6.0 |
| | $1/\gamma$ | U(1, 10) | 0.84 (0.93) | 2.1 (2.0) | 0.93 (0.93) | 2.1 (2.1) | 0.90 | 2.1 |
| BDSS 200-500 tips | $R_0$ | U(1, 5) | 0.94 (0.93) | 1.1 (1.2) | 0.93 (0.93) | 1.2 (1.2) | 0.94 | 1.1 |
| | $1/\gamma$ | U(1, 10) | 0.87 (0.92) | 1.6 (1.7) | 0.88 (0.91) | 1.6 (1.7) | 0.94 | 1.9 |
| | $X_{SS}$ | U(3, 10) | 0.91 (0.90) | 3.5 (3.5) | 0.91 (0.90) | 3.6 (3.6) | 0.97 | 3.5 |
| | $f_{SS}$ | U(0.05, 0.20) | 0.82 (0.78) | 0.087 (0.086) | 0.80 (0.78) | 0.089 (0.089) | 0.94 | 0.110 |
| BD 50-199 tips | $R_0$ | U(1, 5) | 0.90 (0.91) | 1.4 (1.5) | 0.90 (0.90) | 1.4 (1.5) | 0.91 | 1.4 |
| | $1/\gamma$ | U(1, 10) | 0.86 (0.87) | 2.3 (2.2) | 0.85 (0.87) | 2.3 (2.2) | 0.94 | 2.4 |
| BDEI 50-199 tips | $R_0$ | U(1, 5) | 0.93 (0.89) | 1.6 (1.5) | 0.93 (0.89) | 1.6 (1.6) | 0.96 | 1.7 |
| | $1/\varepsilon$ | [0.2, 50] | 0.91 (0.89) | 9.7 (9.3) | 0.88 (0.88) | 10 (9.8) | 0.96 | 12 |
| | $1/\gamma$ | U(1, 10) | 0.95 (0.90) | 2.8 (2.8) | 0.93 (0.90) | 2.9 (2.8) | 0.99 | 3.1 |

For each model and inferred parameter, we compared the different inference methods in terms of 95% confidence interval (95% CI) width and coverage, the latter being defined as the fraction of samples where the true value was within the 95% CI. We compared FFNN-SS and CNN-CBLV methods to BEAST2 on a set of 100 simulations. We did not consider the simulations for which BEAST2 did not converged (2% for BDEI large trees, 5% for BDEI small trees and 15% for BDSS large trees). We evaluated the same metrics on 10,000 simulations for FFNN-SS and CNN-CBLV (in parentheses). For FFNN-SS and CNN-CBLV we performed an approximated parametric bootstrap, while for BEAST2, we considered the entire chain with exception of the initial 10% burn-in. We highlight poor performance (coverage $\leq 0.85$, width $\geq 1/3^{rd}$ of prior) in one of these metrics in red and good performance (coverage $\geq 0.95$) in green. Globally, the 95% CI width and coverage of FFNN-SS and CNN-CBLV are comparable with BEAST2 (while not penalizing BEAST2 for non-converged estimations) and reflect the accuracy of parameter predictions (Fig. 3 and Supplementary Fig. 2). They are slightly better for BDEI with large trees and slightly worse for BDEI with small trees. For details on how the 95% CIs were obtained, see Methods.

**Supplementary Figure 1: Bijectivity of the CBLV representation**

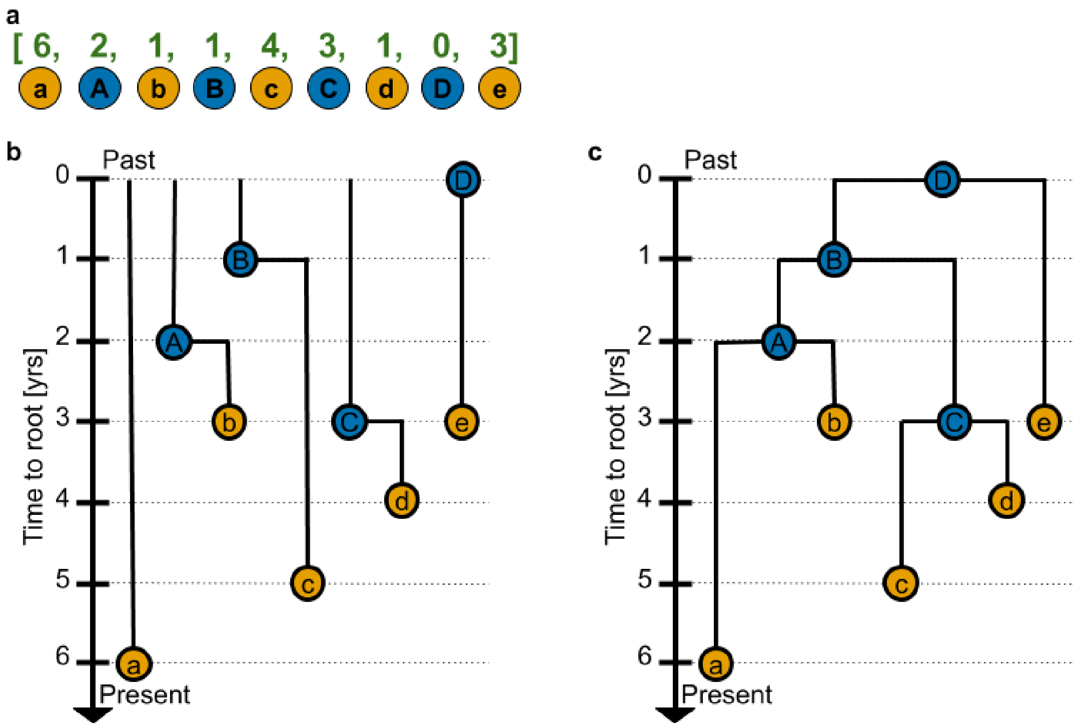


Trees are assumed to be ordered, *e.g.* using ladderization or any other criterion. The inorder tree traversal procedure transforms an ordered tree into a unique vector. We show in this figure using a simple example, how an ordered tree is reconstructed without ambiguity from its vectorial representation, thus demonstrating the bijectivity (1-to-1 correspondence between trees and tree-compatible vectors) of the representation. **a,** shows the Compact Bijective Ladderized Vector (CBLV) representation from Figure 2 a (iii); the nodes are named alphabetically in order of appearance. Lower case letters are highlighted in yellow and represent the external nodes (or tips), upper-case letters are highlighted in blue and represent the internal nodes. **b,** depicts the creation of individual 'paths' comprising each pair of nodes: one external and one internal node, taken directly from the vector representation in the order of appearance. Note that the first external node is not paired with an internal node as it is connected to the root. **c,** shows how the tree reconstruction is achieved, by simply joining the paths from **b,** one by one from left to right. Note that not all vectors correspond to trees, as all entries must be positive or null, plus additional constraints and inequalities (*e.g.*, the second entry (A=2) must be less than (or equal to) the first entry (a=6))**.**

**Supplementary Figure 2: Assessment of deep learning accuracy on small trees.**

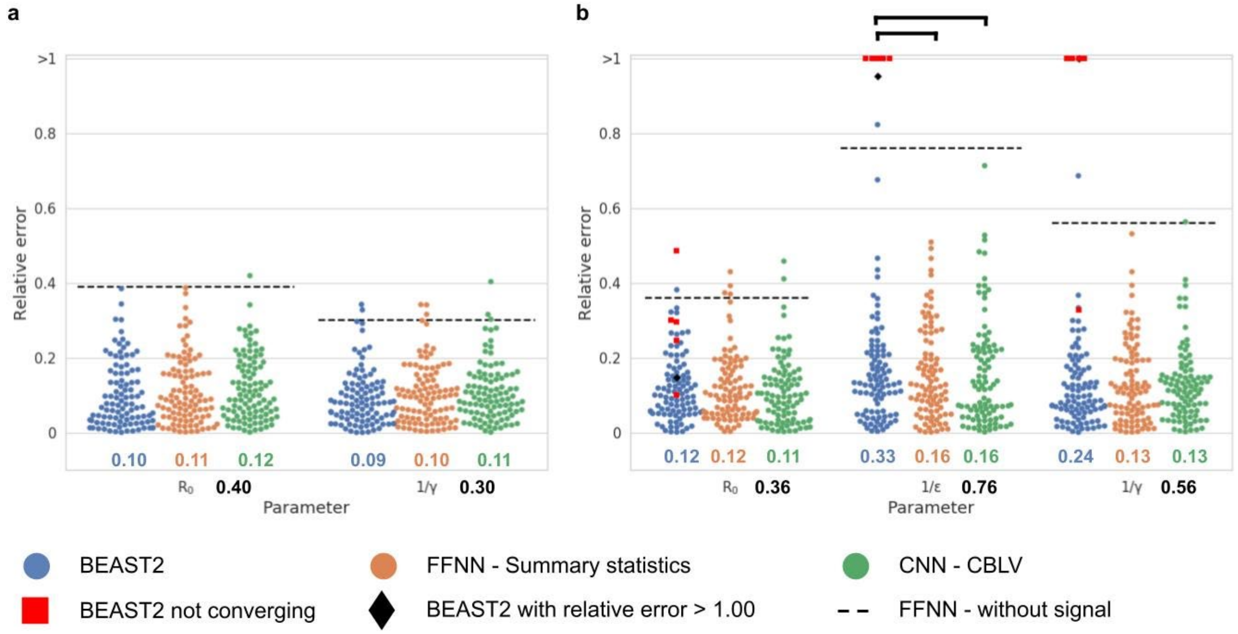


Comparison of inference accuracy by BEAST2 (in blue), FFNN-SS (in orange) and CNN-CBLV (in green) on 100 small test trees (50-199 tips). We compare the relative error for each tree, between the median *a posteriori* estimate by BEAST2 or point estimates by neural networks and the target value for each parameter. We highlight simulations for which BEAST2 did not converge and whose values were thus set to the median of the covered parameter space by depicting them as red squares. We further highlight the analyses with a high relative error (>1.00) for one of the estimates as black diamonds. We compare the relative errors for **a,** BD-simulated, and **b,** BDEI-simulated small trees. Average relative absolute error (MRE) is displayed under each distribution in the corresponding colour. The average error of an FFNN trained on summary statistics but with randomly permuted target is displayed as black dashed line and its value is shown in bold black below the x-axis. The accuracy of the output of each method is compared by paired z-test; $P < 0.05$ is shown as thick full line; non-significant when not shown. The accuracy is similar for the BD model, while the NNs reach better accuracy for BDEI model, while avoiding problems with convergence (5% BEAST2 inferences did not converge).

**Supplementary Figure 3 Accuracy of deep learning methods increases with tree size**

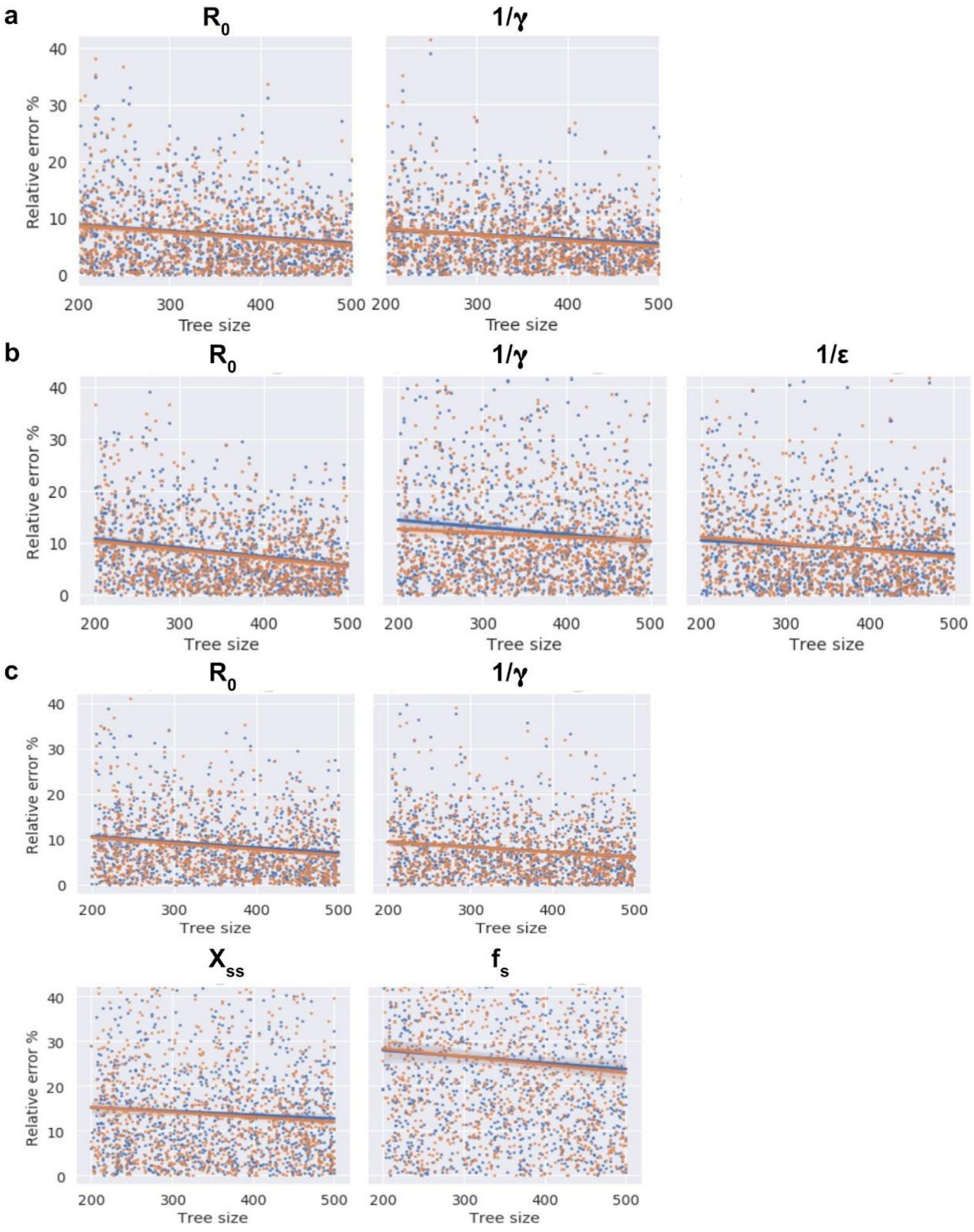


For each model **a,** BD, **b,** BDEI and **c,** BDSS, we display the regression on relative error for each parameter as a function of tree size. We show the error for both CNN-CBLV (in blue) and FFNN-SS (in orange) estimated for 1,000 trees (instead of 10,000 trees for display purposes). As expected, and consistent with statistical learning theory, for each parameter the accuracy increases with tree size. For example, for BDEI, the relative error of $R_0$ is on average 11% for trees of 200 tips and decreases to 6% for trees of 500 tips. Importantly, the relative error of superspreading fraction $f_{SS}$, a parameter that is difficult to estimate, decreases from 28% for trees of 200 tips to 23% for trees of 500 tips.

### Supplementary Figure 4: Assessment of deep learning generalization capabilities

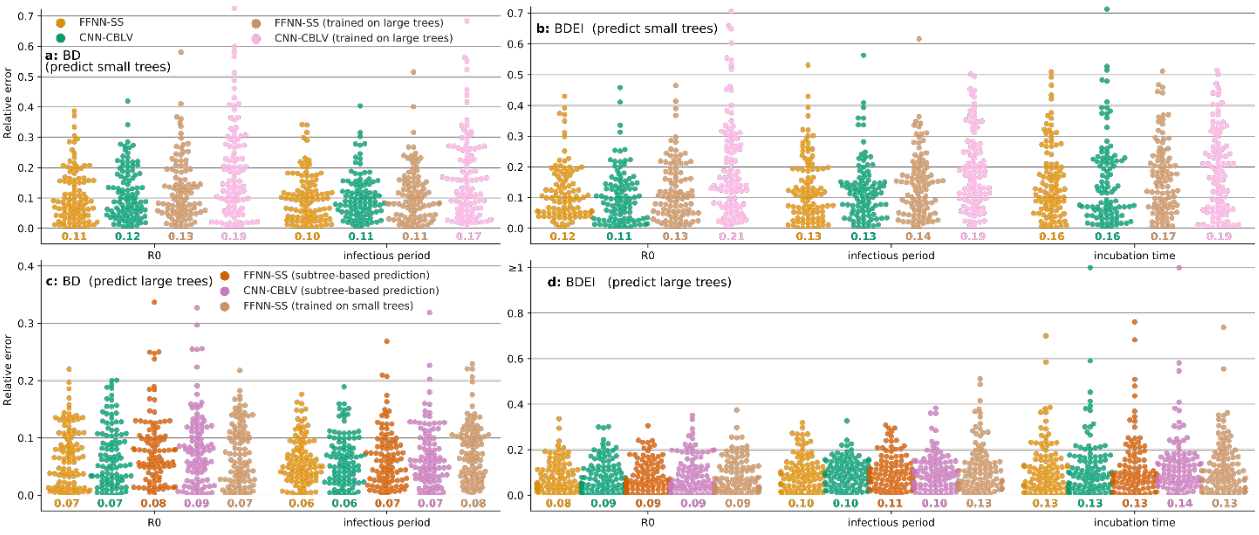


Comparison of inference accuracy by neural networks trained on trees of sizes different from those of the test trees: (top) trained on large trees and evaluated on prediction with small trees (FFNN-SS in beige, CNN-CBLV in pink); and (bottom) trained on small trees and evaluated on prediction with large trees (FFNN-SS in beige, FFNN-SS using subtree picking-and-averaging in red, CBLV-NN using subtree picking-and-averaging in magenta). For comparison, we also show FFNN-SS (orange) and CNN-CBLV (green) trained and evaluated on prediction with trees of compatible sizes (small on top, large on the bottom). The training and testing trees are the same as in **Fig. 3** (large) and **Supplementary Fig. 2** (small). We show the relative error for each test tree. The error is measured as the normalized distance between the point estimates by neural networks and the target value for each parameter. We compare the relative errors for **a, c**, BD-simulated, **b, d**, BDEI-simulated trees. Average relative error is displayed for each parameter and method in corresponding color below each figure.

The results are surprisingly good, especially with summary statistics (FFNN-SS) which are little impacted by these changes of scale as they largely rely on means. Moreover, the subtree picking-and-averaging approach performs well for both FFNN-SS and CNN-CBLV, which confirms the finding in **Fig. 4**.

**Supplementary Figure 5: CNN performs well with CBLV tree representation**

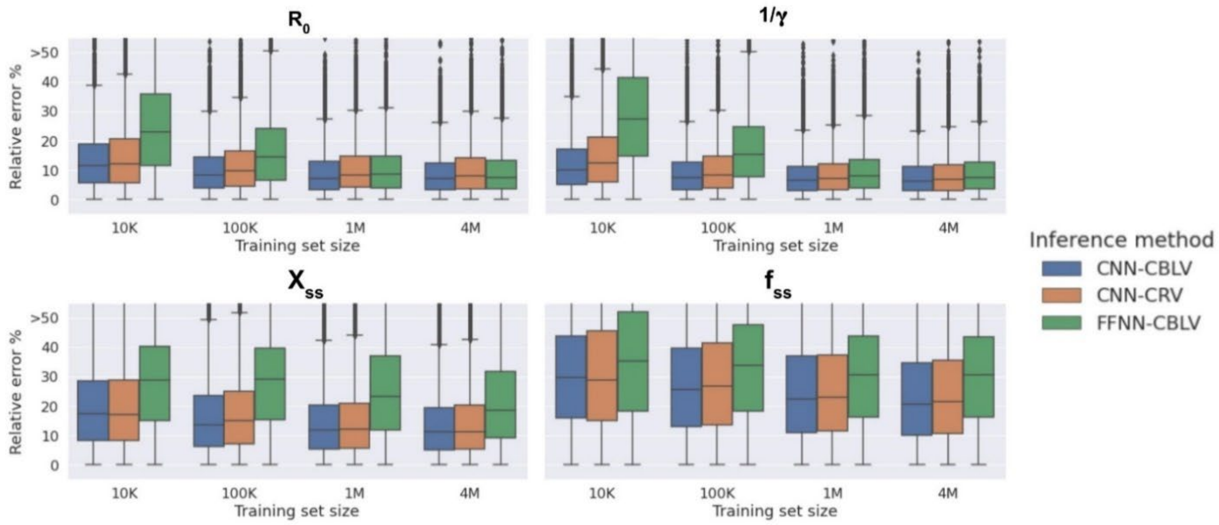


We display percentage absolute error of point estimates from deep learning methods on 10,000 test trees generated with BDSS. We compare CNN-CBLV (in blue) with FFNN-CBLV (in green) and CNN-CRV, which is a CNN trained on a Compact Random Vector (CRV) representation, where all internal nodes are randomly rotated instead of being ladderized, (in orange). In addition, we show the accuracy of these models when trained on varying training set sizes (10K: 10,000; 100K: 1,000,000; 1M: 1,000,000; and 4M: 4,000,000 trees).

For most parameters and training set sizes, the accuracy of CNN-CRV is lower than the one of CNN-CBLV, especially with low number of training examples, while with 4M the accuracy of both methods becomes relatively close. Thanks to ladderization, CBLV is thus enabling the CNN to learn faster and more accurately than if trained on CRV. The difference in performance is even more striking for FFNN-CBLV when compared to CNN-CBLV.CNN-CBLV is more accurate across all parameters and training set sizes, especially for $X_{SS}$ and $f_{SS}$ parameters even after being trained on 4M examples.

**Supplementary Figure 6: How much and how fast FFNN-SS and CNN-CBLV learn?**

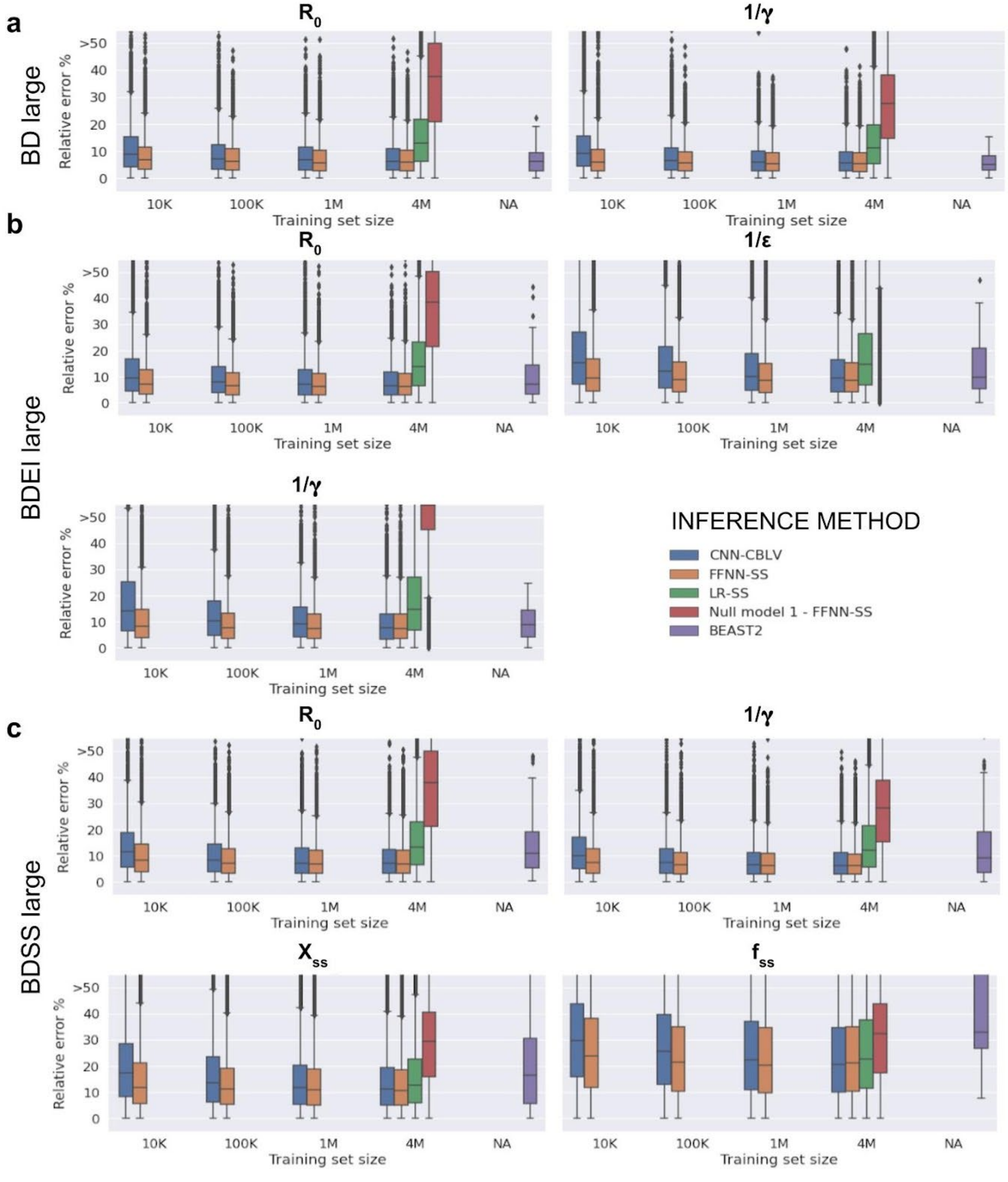


We display the distribution of relative error of individual point estimates from statistical learning methods for 10,000 trees, and median *a posteriori* values from BEAST2 for 100 large trees simulated under each of the three birth-death models **a,** BD, **b,** BDEI, **c,** BDSS. As there are many more points considered for statistical learning methods, the BEAST2 measures are shown for indicative purposes only (in purple). We compare CNN-CBLV (in blue), FFNN-SS (in orange), LR-SS as a baseline model (in green) and "Null model 1", for which the FFNN was trained on summary statistics to predict permuted target values (in red). The Null model maintains the same cost function as other neural networks and thus minimizes the mean percentage relative error in the absence of any signal.

In addition, we show the accuracy of estimated statistical models trained on varying training set sizes (10K: 10,000; 100K: 100,000; 1M: 1,000,000; and 4M: 4,000,000 trees), to study the efficiency of the different learning approaches.

The FFNN-SS accuracy culminates at a training size of 100,000, while CNN-CBLV keeps improving at a training size of 4,000,000. This is consistent with the fact that summary statistics represent high-level information, while the CNN has to learn how to infer parameter values from raw information and thus require many more examples.

The baseline model LR-SS does not reach the same level of accuracy as FFNN-SS and CNN-CBLV for most parameters, but $X_{SS}$ and $f_{SS}$ parameters in BDSS. The relationship between the summary statistics and the studied model parameter values is too complex to be handled by linear regression. Finally, by comparing the accuracies to those of "Null model 1", we show how much information is extracted by the properly set-up deep-learning methods as opposed to a model trained in the absence of signal. In most cases, the accuracy gain is high. The only exception is $f_{SS}$ parameter, where the "Null model 1" has an accuracy of 0.33, while with CNN-CBLV and FFNN-SS we reach an accuracy of around 0.25. These two approaches thus do not estimate this parameter very accurately, most likely because [1] $f_{SS}$ is a fraction of rates and thus cumulates several errors, and [2] the information in the tree on superspreading is low, as the fraction and thus the number of superspreading individuals is low as well. One solution to this problem is to gather more data. Indeed, as shown in Supplementary Fig.7 accuracy increases substantially when learning and inferring on larger trees.

**Supplementary Figure 7: Adding new SS to increase accuracy of FFNN-SS**

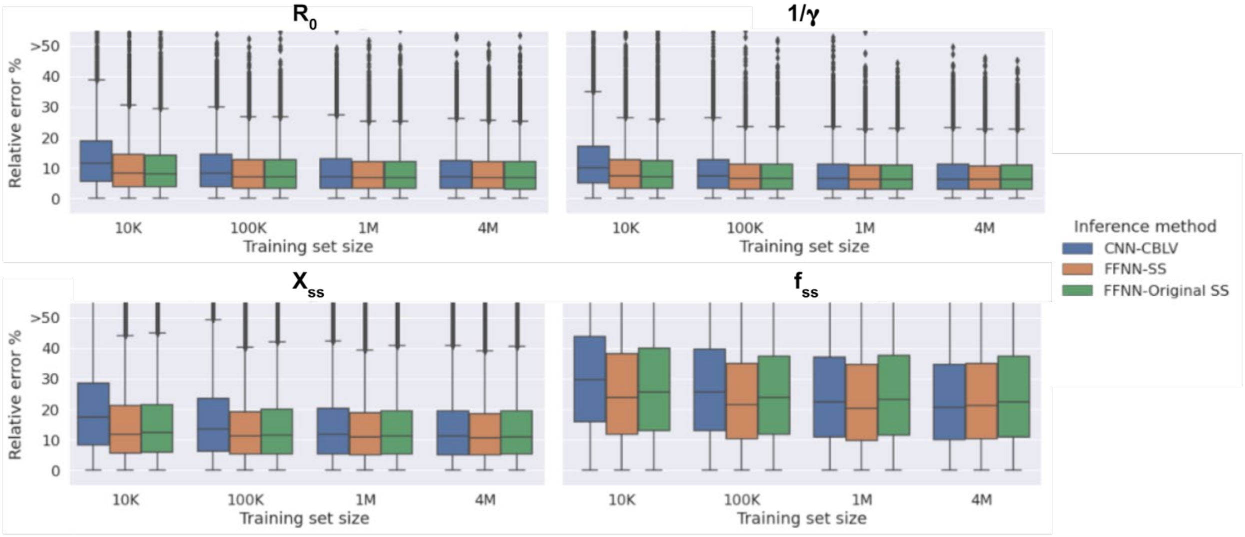


Adding SS on transmission chains improves the accuracy of prediction of superspreading individual frequency $f_{ss}$ in the BDSS model. We display relative absolute error (RE) of point estimates from deep learning methods for 10,000 trees simulated under the BDSS model. We compare CNN trained on CBLV representation with FFNN trained on SS and on Original Saulnier's SS (*i.e.,* without SS on transmission chains), for each parameter of interest. In addition, we show the accuracy of NN models trained on varying training set sizes (10K: 10,000; 100K: 100,000; 1M: 1,000,000; and 4M: 4,000,000 trees). This shows that additional SS enable to decrease the MRE by over 2% for $f_{ss}$ making it comparable to CNN-CBLV accuracy with large training sample (4M).

**Supplementary Figure 8:** ***A priori*** **and** ***a posteriori*** **checks of model adequacy for HIV data**

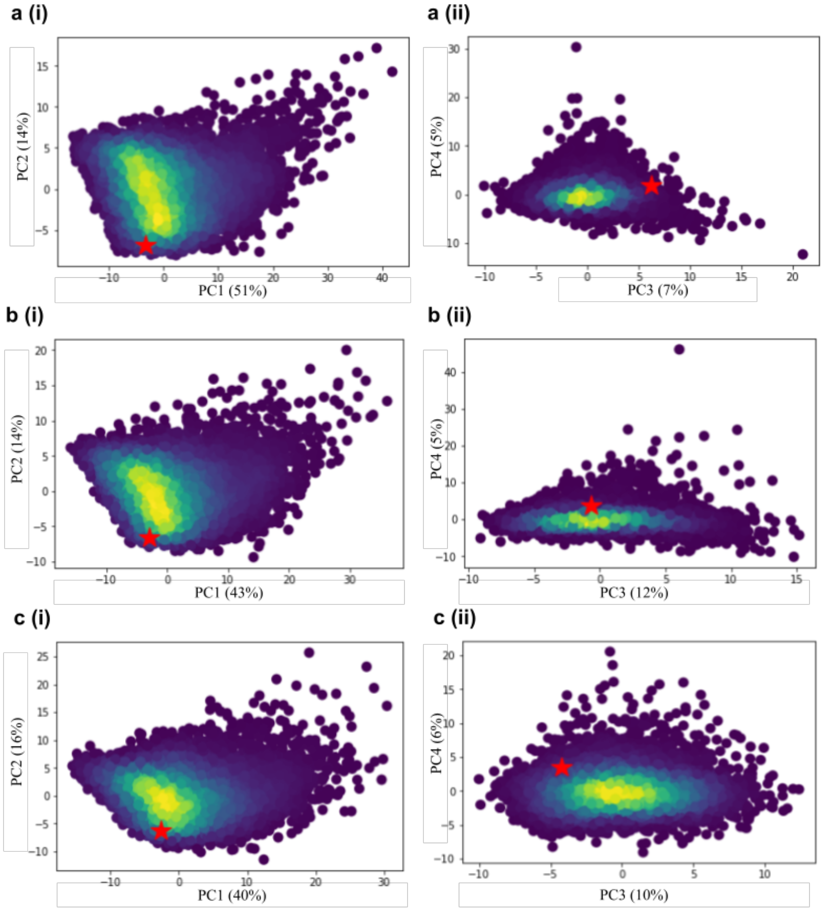


**d**

| Model | # SS outside Min-Max | SS Names |
|---|---|---|
| BD | 5 | ie BL mean 2 and 3, ie BL median 2 and 3, ie BL var 3 |
| BDEI | 5 | min Height, ie BL mean 2 and 3, ie BL median 3, ie BL var 3 |
| BDSS | 0 | - |

**e**

| BDSS | # SS outside Min-Max | SS Names |
|---|---|---|
| FFNN-SS | 13 | max_L, slope_1, mean_s_time, x_16, x_17, x_18, y_3-y_9 |
| CNN-CBLV | 20 | max_L, t_max_L, slope_1, mean_s_time, x_11-x_18, y_2-y_9 |

For each model **a**, BD, **b**, BDEI and **c**, BDSS, we encoded 10,000 simulations from the test set into SS and standardized them. We then performed principal component analysis (PCA) and projected the SS from HIV phylogeny (red star) on these PCA plots. Here we show the projections along **a-c (i)**, the 1st and the 2nd components (PC1 and PC2) and **a-c (ii)**, the 3rd and the 4th components (PC3 and PC4) of the PCA, together with the associated percentage variance explained in parentheses. For each model and projection, the HIV data point is surrounded by the simulations, meaning it resembles globally the simulations and thus we can apply our deep learning (and BEAST2) inference methods under each birth-death model.

Furthermore, we performed more detailed **d**, priori and **e**, posteriori checks using directly the values of summary statistics without a PCA. For each statistics of HIV phylogeny, we checked whether it lays between the minimum and the maximum value of this statistics covered in **d**, test set of 10,000 trees for given model (BD, BDEI or BDSS), **e**, the a posteriori set under BDSS (10.000 simulations, see text). The statistics that were outside the minimum and maximum values are numbered and named in **d** and **e**. All rejected SS in *a posteriori* check (**e**) correspond to the LTT plot (e.g., x and y coordinates), consistent with the fact that the probabilistic, sampling component of the BDSS model is an oversimplification of actual sampling schemes, which depend on contact tracing, sampling campaigns and policies, etc. For details on each statistics, refer to https://doi.org/10.1371/journal.pcbi.1005416.
